# Quantifying cell viability through organelle ratiometric probing

**DOI:** 10.1101/2023.04.26.538448

**Authors:** Rui Chen, Kangqiang Qiu, Guanqun Han, Bidyut Kumar Kundu, Guodong Ding, Yujie Sun, Jiajie Diao

## Abstract

Detecting cell viability is crucial in research involving the precancerous discovery of abnormal cells, the evaluation of treatments, and drug toxicity testing. Although conventional methods afford cumulative results regarding cell viability based on a great number of cells, they do not permit investigating cell viability at the single-cell level. In response, we rationally designed and synthesized a fluorescent probe, PCV-1, to visualize cell viability under the super-resolution technology of structured illumination microscopy. Given its sensitivity to mitochondrial membrane potential and affinity to DNA, PCV-1’s ability to stain mitochondria and nucleoli was observed in live and dead cells, respectively. During cell injury induced by drug treatment, PCV-1’s migration from mitochondria to the nucleolus was dynamically visualized at the single-cell level. By extension, harnessing PCV-1’s excellent photostability and signal-to-noise ratio and by comparing the fluorescence intensity of the two organelles, mitochondria and nucleoli, we developed a powerful analytical assay named *organelle ratiometric probing* (ORP) that we applied to quantitatively analyze and efficiently assess the viability of individual cells, thereby enabling deeper insights into the potential mechanisms of cell death. In ORP analysis with PCV-1, we identified 0.3 as the cutoff point for assessing whether adding a given drug will cause apparent cytotoxicity, which greatly expands the probe’s applicability. To the best of our knowledge, PCV-1 is the first probe to allow visualizing cell death and cell injury under super-resolution imaging, and our proposed analytical assay using it paves the way for quantifying cell viability at the single-cell level.

---

Cells, as the essential unit of organisms, have been broadly studied in relation to biological processes, medical treatments, and nerve conduction. Processes that alter cell viability, including apoptosis, autophagy, and necrosis, are fundamental to eliminating damaged cells and maintaining homeostasis^1,2^. In particular, modulating the specific cellular response of dysregulated cell death, as is prevalent in diseases such as cancer, autoimmune disease, and neurodegeneration^3,4^, has shown success as a therapeutic strategy. Therefore, detecting cell viability is crucial in the discovery of precancerous cells, the evaluation of treatments, and drug toxicity testing^4,5^.

Whereas earlier methods applied to detect cell viability include cellular morphology studies, flow cytometric analysis, and DNA degradation^6,7^, in the past few decades colorimetric assays have been broadly used to monitor cell viability. Among such materials, the well-known yellow 3-(4,5-dimethylthiazol-2-yl)-2,5-diphenyltetrazolium bromide (MTT) is absorbed by live cells and reduced by mitochondrial reductase into purple formazan, which can be quantified by measuring its absorbance using a microplate reader (**Fig. 1a**)^8^. However, because this method can involve factors that may hinder mitochondrial reductase (e.g., amount of cells and incubation time), definite information about cell viability is difficult to obtain^9^. Moreover, owing to its high susceptibility to metabolic interference, MTT-based assay can easily provide false-positive results^4^. Such assays also report the cumulative results of cell viability based on a great number of cells and do not permit investigating cell viability at the single-cell level. Consequently, next-generation biological tools are urgently needed to monitor cell viability on individual cells and thereby enable the rapid screening of drug treatment, minimize experimental error due to external interference, and reduce medical costs.

**Fig. 1.**
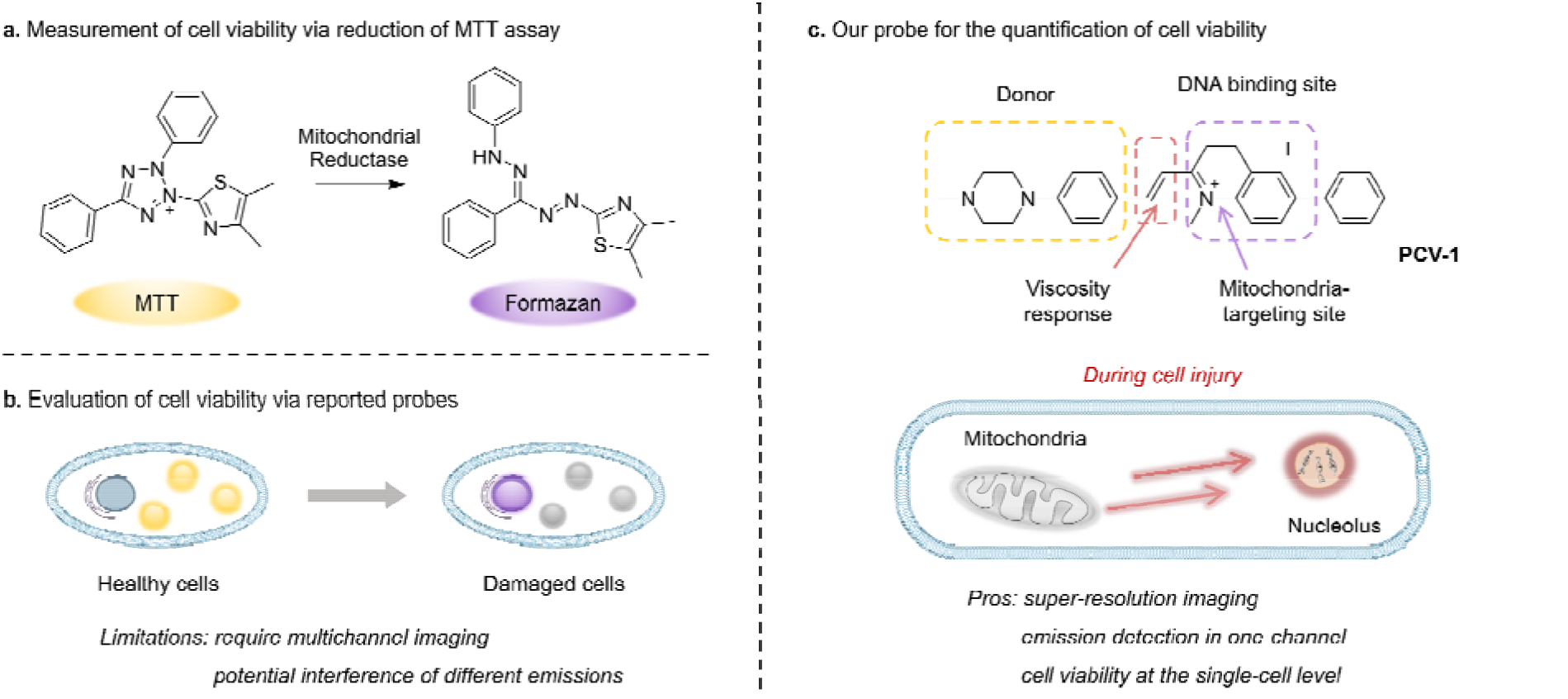
Schematic illustration for assessing cell viability. **a** Measurement of cell viability via the conventional MTT assay. **b** Evaluation of cell viability via previously reported probes. **c** The design strategy of PCV-1 and a schematic illustration for quantifying cell viability at the single-cell level.

While fluorescence microscopy allows the in situ, real-time observation of subcellular organelles and biological processes, fluorescence imaging, as a non-invasive technique that causes relatively little damage, readily accommodates the study of cell viability. Therefore, several fluorescent probes have been developed to distinguish live from dead cells based on various tracking mechanisms and are thus versatile tools for biological research and applications in medicine^5^. As one such fluorescent dye, 4’,6-diamidino-2-phenylindole (DAPI) is widely used to visualize nuclei in dead cells due to its weak ability to penetrate intact plasma membranes in live cells.

In past research, mitochondrial membrane potential (ΔΨ_m_), as a vital feature representing mitochondrial function, has been used as an indicator to discriminate the states of cells^10,11^. A decrease in mitochondrial membrane potential signifies a disturbance in mitochondrial homeostasis that can lead to cell dysfunction or even death. Tian et al., for instance, reported a dual-color fluorescent probe that migrated from mitochondria to the nucleus when the mitochondrial membrane potential decreased during apoptosis; in that way, cell viability could be monitored by identifying the localization and fluorescent intensity of the probe (**Fig. 1b**)^12^. However, the dual-color probe demonstrated several limitations, including the need for a multichannel fluorescence microscope, the potential overlap and interference of different emissions, and the incomparability of fluorescence images from various channels. Meanwhile, Li et al. have developed a red fluorescent probe that can reflect different situations of mitochondrial membrane potential (i.e., normal, decreasing, and vanishing) according to various fluorescence morphologies of the nucleus and mitochondria and thereby indicate the cell’s state^13^. However, the probe’s main limitation is that cell viability cannot be quantitatively assessed by analyzing images owing to the high degree of fluorescence interference between the two organelles and the poor image quality available with conventional confocal microscopy. Thus, clearly observing the interaction of organelles and efficiently quantifying cell viability using fluorescence microscopy remains challenging, especially at the single-cell level.

Recently, super-resolution imaging, as a cutting-edge technology able to overcome the Abbe’s diffraction limit (<200 nm) of conventional confocal microscopy has been greatly applied to observe subcellular organelles and biological interactions^14-18^. Therefore, fluorescence microscopy with super-resolution imaging might be ideal for synchronously visualizing the dynamics of organelles and consequently quantifying cell viability. In view of the size of eukaryotic organelles, including lysosomes (i.e., 100–1,200 nm), mitochondria (i.e., 500–5,000 nm), and nuclei (i.e., approx. 10 μm)^19^, structured illumination microscopy (SIM) may be a promising alternative, especially given its super spatial resolution (i.e., approx. 100 nm), short acquisition time, and only slight photodamage^20-23^.

Against this backdrop, we rationally designed and synthesized a fluorescent probe PCV-1 to visualize cell viability under SIM. As shown in **Fig. 1c**, PCV-1 consists of a donor–π–acceptor structure, in which the positively charged π-conjugated system endows PCV-1 with the ability to target mitochondria given the highly negative mitochondrial membrane potential in healthy cells^24,25^. The cationic methylquinolinium moiety gives PCV-1 a high affinity to DNA due to their electrostatic interaction^26,27^. PCV-1 exhibited extended π-conjugation but decreased π-π stacking compared to the cell viability probe previously reported, based on its molecular design^12^. Moreover, owing to its alkene bridge, PCV-1 can readily respond to viscosity^28,29^. In an environment of high viscosity, PCV-1’s molecular rotation is likely restricted, thereby rendering a more planar configuration that reduces its non-radiative decay and hence increases the emission intensity. By performing the fluorescence imaging of PCV-1-stained HeLa cells using SIM, we observed mitochondria and nucleoli as distinct organelles in live and dead cells in single-channel mode, respectively. Moreover, owing to the excellent super-resolution imaging and viscosity-enhanced emission, PCV-1 is able to provide more subcellular details that cannot be achieved by conventional confocal microscopy. Thus, during cell injury induced by drug treatment, PCV-1’s migration from mitochondria to the nucleolus was dynamically visualized at the single-cell level. Beyond that, based on PCV-1’s translocation from mitochondria to the nucleolus following damage and by comparing the two organelles’ fluorescence intensity at the subcellular level, we developed a new analytical assay, *organelle ratiometric probing* (ORP), that can be used to quantitatively analyze and efficiently assess the viability of individual cells. In the ORP analysis with PCV-1, we systematically investigated the relationship between the various fluorescence intensity ratios of organelles and their corresponding cell viability values at different stages, as a result, identified 0.3 as the cutoff point for assessing whether adding a drug will cause apparent cytotoxicity. Furthermore, regardless of various routes to cell death, including apoptosis, autophagy, and necrosis, a ratio significantly higher than 0.3 for the ORP analysis with PCV-1 indicated nearly dead cells. To our knowledge, PCV-1 is the first probe to allow visualizing cell death and cell injury under super-resolution imaging, and our proposed ORP assay using PCV-1 enables the quantification of cell viability at the single-cell level.

## Results and discussion

### Synthesis, photophysical properties, and computational calculations of PCV-1

As shown in **Fig. 2a**, the synthesis of PCV-1 began with commercially available 6-bromo-2-methylquinoline. The intermediate 2-methyl-6-phenylquinoline was generated by a typical Suzuki coupling reaction between 6-bromo-2-methylquinoline and phenylboronic acid. Next, methylation at the nitrogen position of 2-methyl-6-phenylquinoline using methyl iodide resulted in 1,2-dimethyl-6-phenylquinolin-1-ium. Meanwhile, 1-methylpiperazine reacting with *p*-fluorobenzaldehyde under an alkaline conditions formed another intermediate 4-(4-methylpiperazin-1-yl) benzaldehyde with a nearly quantitative yield. Finally, the Knoevenagel condensation reaction between 1,2-dimethyl-6-phenylquinolin-1-ium and 4-(4-methylpiperazin-1-yl) benzaldehyde in anhydrous ethanol generated our designed product, PCV-1. Details of the synthesis procedure and characterizations of ^1^H, ^13^C NMR, and MS for intermediates and products are included in the Supplementary Information (**Supplementary Fig. 1–5**).

**Fig. 2.**
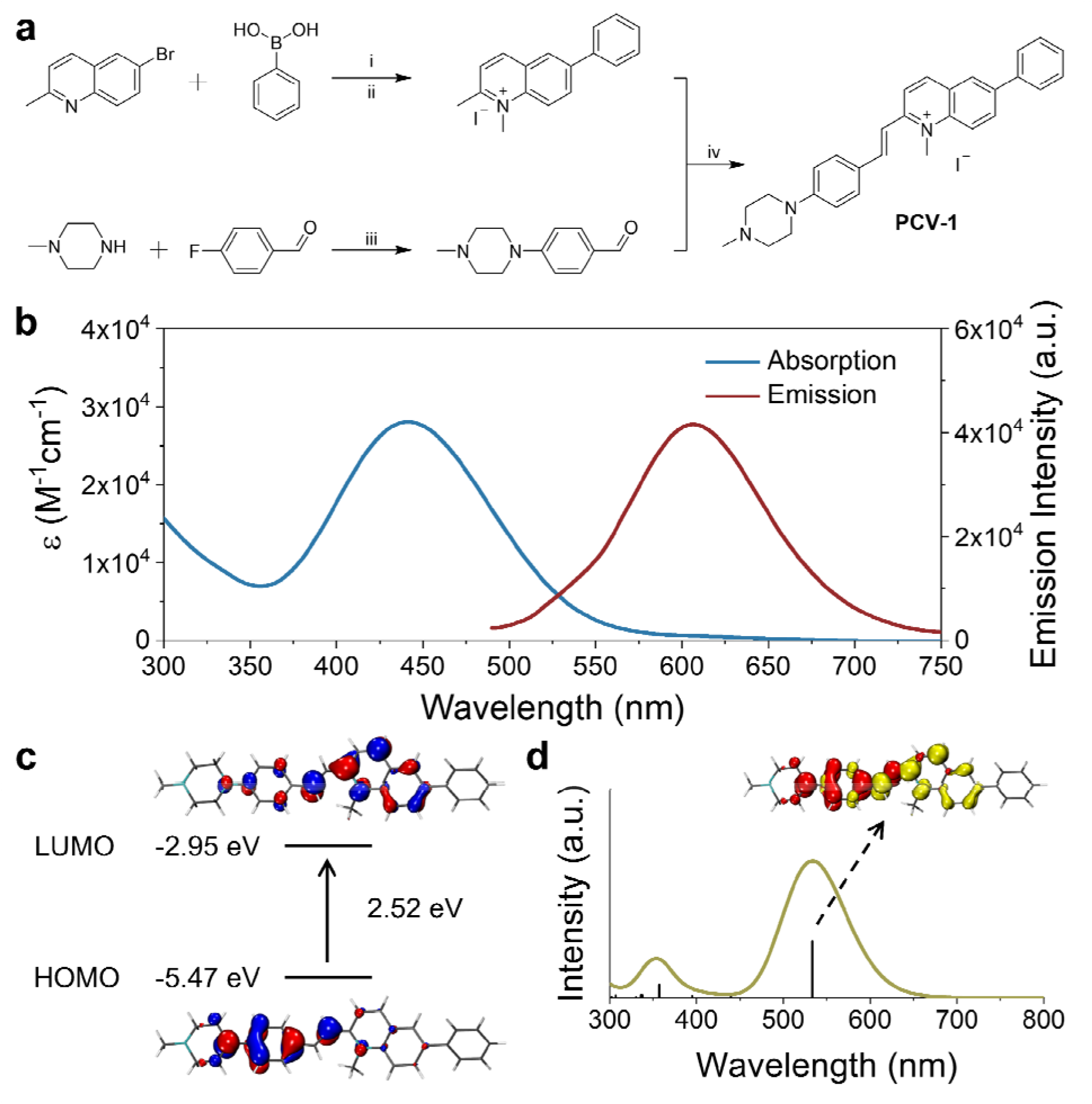
Synthesis, photophysical properties, and computational calculations of PCV-1. **a** Synthetic scheme of the designed fluorescent probe PCV-1. (i) Pd(PPh_3_)_4_, K_2_CO_3_, Bu_4_NBr, toluene, EtOH, H_2_O, 100 °C; (ii) CH_3_I, EtOH or CH_3_CN, 80 °C; (iii) K_2_CO_3_, DMF, 100 °C; (iv) EtOH, 80 °C. **b** Absorption and emission spectra of PCV-1. **c** Calculated HOMO and LUMO of PCV-1. **d** Calculated absorption spectrum of PCV-1 and EDDM of the 1^st^ singlet excited state of PCV-1 (red and yellow color indicate decrease and increase in electron density, respectively).

With the designed probe PCV-1 in hand, we first investigated its photophysical properties in aqueous solution. As shown in **Fig. 2b**, PCV-1 exhibited a broad absorption spectrum with a tail exceeding 570 nm due to its large π–π conjugation. The maximum absorption of PCV-1 was located at 442 nm with a high molar extinction coefficient of 28,060 M^−1^cm^−1^. Under 450 nm excitation, PCV-1 showed a strong red emission with a maximum peak at 606 nm, which is favorable for bioimaging applications. The Stokes shift of PCV-1 was beyond 160 nm due to its charge-enhanced intramolecular charge transfer system and extended conjugation length^30^. Moreover, when PCV-1’s emission spectra were investigated in different pH buffer solutions, no apparent change in emission intensity was observed in the pH range of 4.0–8.0, as shown in **Supplementary Fig. 6**. Next, the potential influence of biochemical species on PCV-1’s emission intensity was investigated. As illustrated in **Supplementary Fig. 7**, various sources of interference in the cytoplasm and microenvironment were added to the aqueous solution of PCV-1, including the cations of Na^+^, K^+^, Zn^2+^, Fe^2+^, Fe^3+^, Mg^2+^, Ag^+^, and Ni^2+^, the anions of NO_3_^−^, SO_4_^2−^, AcO^−^, HCO_3_^−^, HSO_3_^−^, SCN^−^, F^−^, Cl^−^, Br^−^, and I^−^, the biomolecules of GSH, Hcy, and Cys, and the reactive substance H_2_O_2_. No evident changes in PCV-1’s emission intensity were observed in the presence of any aforementioned species.

To shed light on PCV-1’s electronic transitions, density functional theory (DFT) calculations were performed using the Gaussian 16W program package (**Supplementary Table 1, 2**). As shown in **Fig. 2c**, PCV-1’s highest occupied molecular orbital (HOMO) was located in the electron-donating phenyl piperazine group. With the introduction of the positive charge via methylation, the central methylquinolinium unit showed greater electronegativity. As expected, PCV-1’s lowest unoccupied molecular orbital (LUMO) was centered on the electron-accepting methylquinolinium moiety. The calculated energies of HOMO and LUMO were −5.47 eV and −2.95 eV, respectively, resulting in an energy gap of 2.52 eV. Moreover, time-dependent DFT calculations indicated that PCV-1’s lowest singlet excited state S_1_ was mainly HOMO to LUMO transition (see the electron density difference map (EDDM) image in Fig. 2d) and located at 533 nm (**Fig. 2d**), which generally matched the experimental absorption spectrum (**Fig. 2b**).

### Viscosity-enhanced optical properties of PCV-1

The viscosity of subcellular organelles is far higher than cytosol’s^31-33^. Lysosomes in particular are known to contain an internal environment with viscosity ranging from 50 to 90 cP^34^. The average viscosity of mitochondria, by comparison, is estimated to be approximately 62 cP^35^. Therefore, the large difference in viscosity between cell cytosol and organelles can facilitate subcellular bioimaging that results in high brightness, less photodamage, and a large signal-to-noise ratio.

Fluorescent probes designed with rotatable C_sp_^2^–C_sp_^2^ bonds have recently been developed to respond to the viscosity of organelles^36,37^. Because PCV-1 is composed of a donor–π–acceptor structure, rotatable C_sp_^2^–C_sp_^2^ bonds also exist between the vinylene group and the electron-donating and -accepting moieties, as shown in **Fig. 3a**. To investigate PCV-1’s optical properties in response to viscosity, we collected PCV-1’s absorption and emission spectra in water upon the addition of glycerol. Because PCV-1’s molecular rotation is likely restricted in solutions of high viscosity, PCV-1 forms a more planar configuration with better conjugation. As a result, PCV-1’s absorption peak wavelength showed a 32-nm redshift in glycerol solution compared with that in water (**Fig. 3b**). Next, we measured PCV-1’s emission spectra in solutions with viscosities ranging from 0.9 to 945 cP. As shown in **Fig. 3c**, PCV-1 initially exhibited relatively weak emission intensity in solutions of low viscosity. However, as the viscosity increased, the molecular rotation is restricted and the excited state energy was primarily released through radiative decay^28^. Therefore, PCV-1 presented a significantly enhanced (i.e., 187-fold) emission intensity in glycerol. By fitting via the F□rster–Hoffmann equation, as shown in the inset of **Fig. 3c**^37,38^, PCV-1’s emission intensity demonstrated an excellent linear relationship (*R*^2^ = 0.98) with viscosity. Meanwhile, its excited-state lifetime was independent of concentration and absorption^39^. As shown in **Fig. 3d**, PCV-1’s excited-state lifetime in glycerol was much longer than that in water, which further confirms the different molecular rotation and π-conjugation of PCV-1 in solutions of high and low viscosity.

**Fig. 3.**
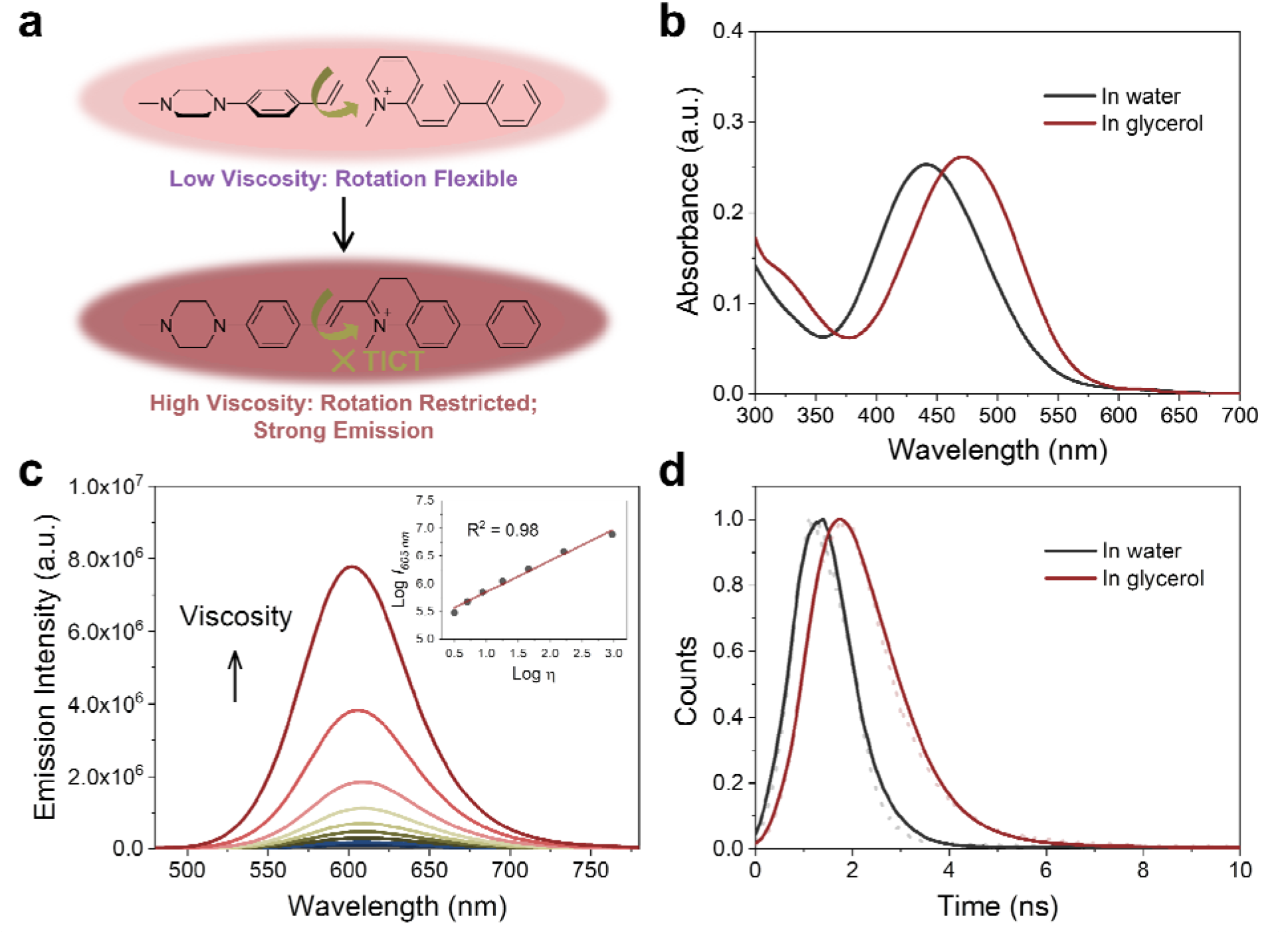
Viscosity-enhanced optical properties of PCV-1. **a** Responsive mechanism of PCV-1 towards viscosity. **b** Absorption spectra of PCV-1 in water and glycerol. **c** Emission spectra of PCV-1 in water upon the addition of glycerol (λ*_ex_* = 450 nm). Inset: scatter plots and linear relationship between Log *I_605 nm_* and Log η for PCV-1. **d** Emission decay plots of PCV-1 in water and glycerol.

To examine whether PCV-1’s strong emission could also be attributed to the polarity of solvents, PCV-1’s emission spectra were collected in different solvents. As illustrated in **Supplementary Fig. 8a**, PCV-1 showed significantly high emission intensity only in glycerol. When it was dissolved in solvents of different polarity with low viscosity, the observed emission intensity was nearly negligible compared with that in glycerol (**Supplementary Fig. 8b**). Such an infinitesimal polarity effect makes PCV-1 a highly viscosity-sensitive probe in complex biological microenvironments. Moreover, PCV-1’s emission intensity remained unchanged after 300 s of irradiation in both low- and high-viscosity solutions (**Supplementary Fig. 9**), which confirms its superior photostability. Taken together, our photophysical results strongly suggest that PCV-1 is a promising viscosity-responsive probe that is well-suited for imaging highly viscous organelles, such as mitochondria.

### Specific localization of PCV-1 in cells

Encouraged by PCV-1’s outstanding photophysical properties, we next explored its application in bioimaging. First, PCV-1’s toxicity was evaluated using the colorimetric cell viability assay with a Cell Counting Kit-8 (CCK-8). The CCK-8 assay was performed with the highly water-soluble tetrazolium salt WST-8, which can be reduced by dehydrogenase in healthy cells and generate yellow WST-8 formazan. Per those results, the higher activity of dehydrogenase implies greater cell viability. The CCK-8 assay results indicated that PCV-1 had no effect on cell viability at various concentrations after 24 h treatments (**Supplementary Fig. 10**). Next, we performed the fluorescence imaging of PCV-1 via SIM. As depicted in **Fig. 4a**, the SIM images of PCV-1-stained Hela cells showed filamentous fluorescence that might indicate mitochondria.

**Fig. 4.**
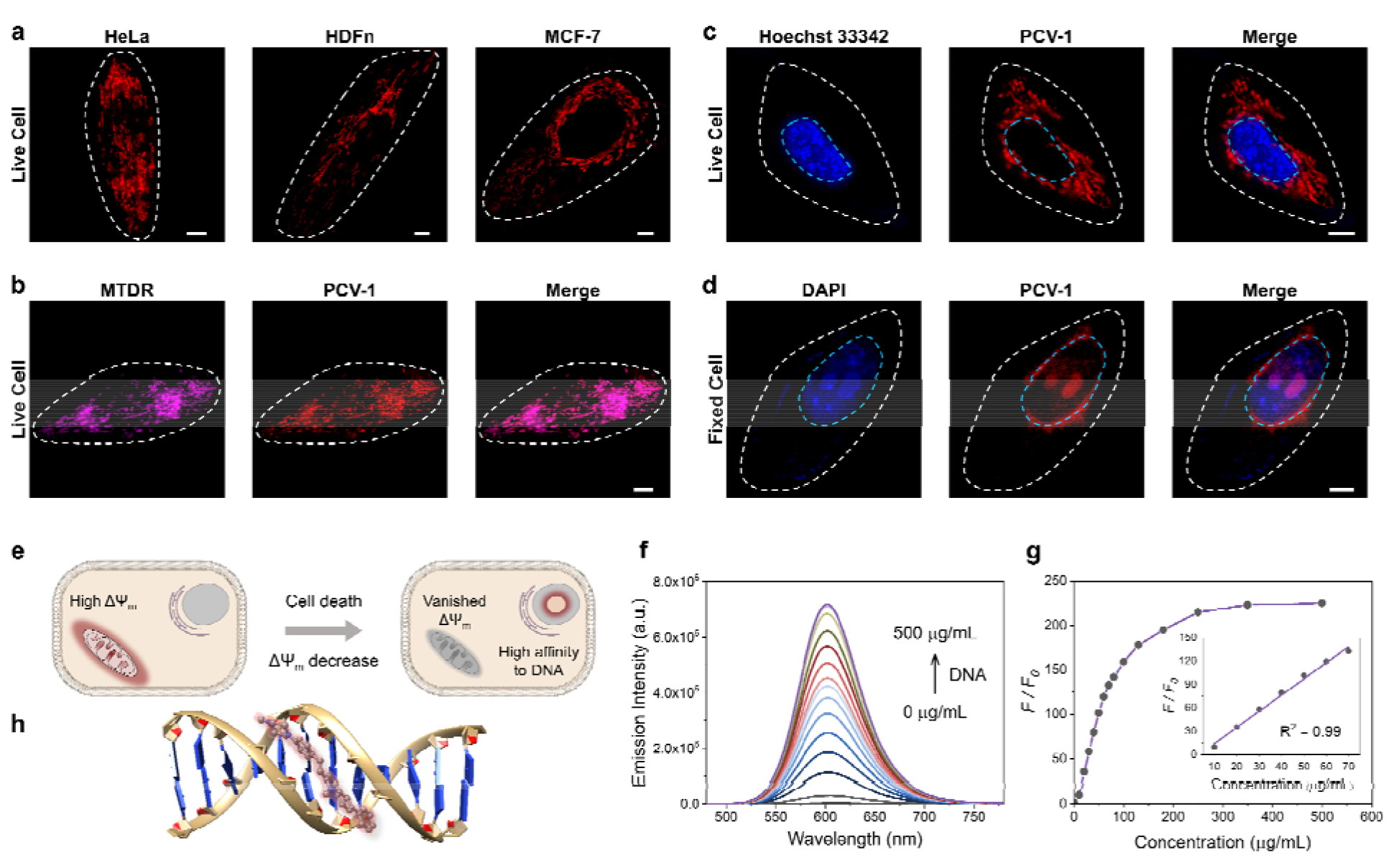
Specific localization of PCV-1 in cells. **a** SIM images of PCV-1-stained live HeLa, HDFn, and MCF-7 cells. **b** SIM images of live HeLa cells co-stained with MTDR and PCV-1. **c** SIM images of live HeLa cells co-stained with Hoechst 33342 and PCV-1. **d** SIM images of fixed HeLa cells co-stained with DAPI and PCV-1. PCV-1 channel: Ex 561 nm, Em 570-640 nm; MTDR channel: Ex 640 nm, Em 660-735 nm; Hoechst 33342 and DAPI channels: Ex 405 nm, Em 425-475 nm; scale bars: 5 μm. **e** Schematic illustration for different targeting abilities of PCV-1 in live and dead cells. **f** Fluorescence titration experiments of PCV-1 in buffer solution with increasing the concentration of calf thymus DNA (0-500 μg/mL), λ*_ex_* = 450 nm. **g** Plots of emission intensity enhancements at 600 nm versus concentration of calf thymus DNA. Inset: scatter plots and linear relationship between emission intensity enhancements and DNA concentration. **h** Binding mode of PCV-1 to DNA via molecular docking calculations.

The same filamentous fluorescence images were also observed in other cell lines, including fibroblast cells (i.e., HDFn) and breast cancer cells (i.e., MCF-7). Given those observations, co-localization experiments of PCV-1 were conducted with the commercial mitochondria dye MitoTracker Deep Red (MTDR, **Fig. 4b**), and the lysosome dye LysoTracker Deep Red (LTDR, **Supplementary Fig. 11**). The fluorescence images of PCV-1 showed a higher overlap with MTDR (Pearson’s correlation coefficient (PCC) = 0.91, **Supplementary Fig. 12a**) than LTDR (PCC = 0.34, **Supplementary Fig. 12b**), which indicates that PCV-1 can be used as a mitochondrion-targeting probe. To further gauge PCV-1’s ability to localize in mitochondria, we investigated its lipophilicity by calculating the *n*-octanol–water partition coefficient value (i.e., logP). As shown in **Supplementary Fig. 13**, the logP value of PCV-1 was 0.7, thereby indicating that PCV-1 has high lipophilicity. According to Horobin et al.’s theory, a cation with a logP value from 0 to 5 has a high probability of mitochondrial localization^40-42^. Thus, the co-localization images together with the logP value demonstrate the mitochondrial localization of PCV-1 in live cells.

Afterwards, when we investigated the fluorescence imaging of PCV-1 after drug treatment, we found that PCV-1’s fluorescence morphology in dead cells completely differed from that in live cells. As shown in **Supplementary Fig. 14**, no filamentous fluorescence was presented in the cytoplasm region of the dead cells. Instead, PCV-1 was found to gather in the center of cells, where it exhibited elliptical morphology and high fluorescence intensity. Based on these results, we speculated that the new fluorescent region was located in the nucleus and PCV-1 escaped from mitochondria to the nucleus during cell death. To test this hypothesis, we subsequently co-stained it with the nuclei dyes of live cells (i.e., Hoechst 33342) and dead cells (i.e., DAPI). As shown in **Fig. 4c**, no evident overlap between the blue channel of Hoechst 33342 and the red channel of PCV-1 was observed, which confirmed that PCV-1 was only localized in mitochondria in live cells. In sharp contrast, PCV-1 could also cross the nuclear envelope and displayed strong red fluorescence in fixed cells, which highly overlapped with the blue fluorescence from DAPI, suggesting that PCV-1 can selectively stain the nuclei in dead cells (**Fig. 4d**). Moreover, the strong red emission of PCV-1 was greatly concentrated in the central nuclear region with spherical morphologies, which indicated that PCV-1 was primarily distributed in the nucleolus^43^. Thus, we suspected that the different distributions of PCV-1 in live and dead cells were impacted by the mitochondrial membrane potential. As depicted in **Fig. 4e**, PCV-1 was abundantly enriched in mitochondria due to the cationic quinolinium group in PCV-1 and the highly negative mitochondrial membrane potential in healthy cells. However, in dead cells no longer with mitochondrial membrane potential, PCV-1 targeted the nucleolus via electrostatic attraction and possible intercalation.

To understand the mechanism by which PCV-1 targets the nucleolus in dead cells, we performed a DNA titration experiment using calf thymus DNA (ctDNA). As shown in **Supplementary Fig. 15**, as the concentration of ctDNA in the PCV-1 buffer solution increased, the absorption spectra of PCV-1 showed a clear redshift indicating that PCV-1’s π-conjugation has been expanded. Meanwhile, the emission intensity of PCV-1 gradually enhanced with DNA binding, as illustrated in **Fig. 4f**. When the concentration of ctDNA reached 500 μg/mL, a 225-fold enhancement of PCV-1’s emission intensity was observed (**Fig. 4g**). In addition, PCV-1’s enhanced emission intensity demonstrated an exceptional linear relationship (*R*^2^ = 0.99), with the concentration of ctDNA in the range of 10–70 μg/mL. Based on the properties of PCV-1 in response to viscosity described before, we thus surmise that the light-on behavior of PCV-1 in the presence of DNA can also be due to the restriction of molecular rotation. Therefore, we conducted molecular docking studies between DNA and PCV-1 using the AutoDock Vina software^44^. The results of the docking simulation suggested that PCV-1 was strongly bound to the minor grooves of DNA (**Fig. 4h**). The calculated binding energy was −10.5 kcal mol^−1^, which is comparable to that of the commercial Hoechst 33342 (−12.2 kcal mol^−1^) and better than that of a previously reported nuclei probe^12^. Overall, the aforementioned imaging, emission titration, and molecular docking results collectively demonstrated that PCV-1 can differentiate live and dead cells via the distinct fluorescence morphologies and intensities of different organelles.

### PCV-1’s excellence in super-resolution imaging

To acquire better fluorescence images at the single-cell level as a means to quantify cell viability, we explored PCV-1’s imaging properties in super-resolution imaging. We first compared the fluorescence images of PCV-1 in HeLa cells captured by conventional confocal microscopy and SIM. Confocal imaging exhibited the high spatial overlap of fluorescence due to its low resolution (**Fig. 5a**), while SIM imaging afforded superior imaging quality, with clearly filamentous mitochondria fluorescence and less background fluorescence (**Fig. 5b**). Benefiting from the state-of-the-art SIM technology, PCV-1 can visualize previously invisible subcellular details in biological systems.

**Fig. 5.**
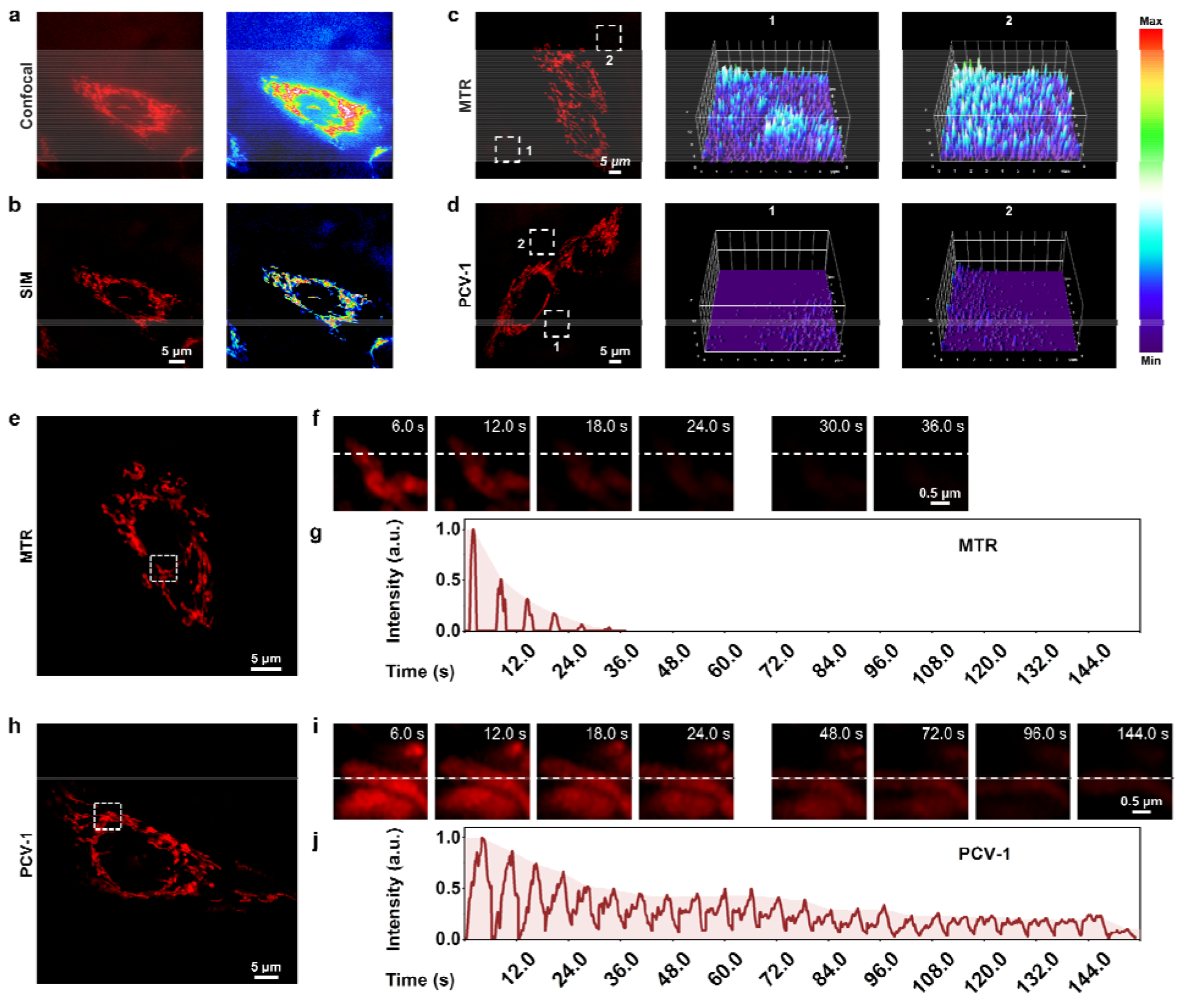
PCV-1’s excellence in super-resolution imaging. **a** Confocal image of HeLa cell stained with PCV-1 and corresponding fluorescence intensity distribution. **b** SIM image of HeLa cells stained with PCV-1 and corresponding fluorescence intensity distribution. SIM images of HeLa cells labeled with **c** MTR or **d** PCV-1 and corresponding 3D fluorescence intensity plots of rectangles of integration, constructed by ImageJ. **e, f** Photobleaching property of MTR under continuous 561 nm laser irradiation at 100% power mode. **g** Normalized fluorescence intensity changes of MTR on the white dotted line in **f**. **h, i** Photobleaching property of PCV-1 under continuous 561 nm laser irradiation at 100% power mode. **j** Normalized fluorescence intensity changes of PCV-1 on the white dotted line in **i**. PCV-1 channel: Ex 561 nm, Em 570-640 nm; MTR channel: Ex 561 nm, Em 570-640 nm.

As indicated, PCV-1 showed a large Stokes shift and viscosity-enhanced fluorescence intensity (**Fig. 2b and 3c**), which can effectively overcome the background noise created by excitation light and increase the signal-to-noise ratio because of high viscosity in mitochondria. Therefore, we chose the commercial mitochondria dye MitoTracker Red (MTR) for comparison to confirm these advantages. The SIM images of HeLa cells labeled with MTR and PCV-1 were taken in the same condition. We randomly picked two regions of background in MTR- and PCV-1-stained SIM images and plotted their corresponding 3D fluorescence intensity distribution maps, as illustrated in **Fig. 5c** and **5d**. The results elucidated that the background fluorescence of PCV-1-stained SIM images was significantly lower than that of MTR-stained SIM images. Beyond that, PCV-1 showed a higher average fluorescence intensity in mitochondria than using MTR (**Supplementary Fig. 16**). In view of these results, PCV-1 can be expected to effectively improve the signal-to-noise ratio of fluorescence imaging and thus facilitate the study of hyperfine structures of cells.

Resistance to photobleaching is a vital ability of probes that determines whether it can function for long-time fluorescence tracking^22,23^. According to the chemical structure and previous fluorescence stability tests, PCV-1 seems to be endowed with a high resistance to photobleaching. To confirm its stability for imaging applications, we performed a photobleaching comparison experiment between MTR and PCV-1 in HeLa cells (**Fig. 5e, h**). Under 100% laser power mode with the same settings, the fluorescence of MTR became rapidly bleached (**Fig. 5f**), while PCV-1 exhibited high resistance to photobleaching (**Fig. 5i**). Specifically, as shown in **Fig. 5g, j**, the photobleaching lifetime of PCV-1 was more than 144 s, or 4 times longer than that of MTR (i.e., 36 s). Furthermore, when the live HeLa cells stained with MTR and PCV-1 were incubated for 24 h for observation (**Supplementary Fig. 17**), both types of cells exhibited fluorescence in mitochondria. On the following day, however, the fluorescence of mitochondria stained with MTR completely disappeared, whereas PCV-1 still showed strong filamentous fluorescence in mitochondria. Thus, we conclude that PCV-1 has superior photostability in live cells.

With all of the outstanding imaging properties described above, including exceptional super-resolution imaging, high signal-to-noise ratio, and great resistance to photobleaching, PCV-1 is expected to be an excellent biological tool for visualizing interactions between organelles at the single-cell level.

### Quantifying cell viability using the ORP analysis with PCV-1

According to the above results, PCV-1 is located in the mitochondria of live cells but escapes to the nucleolus when cells die. Thus, we speculated that there might be a state in which PCV-1 targets both mitochondria and the nucleolus during cell injury. To track the fluorescence morphologies of PCV-1 in the process of cell injury, carbonyl cyanide *m*-chlorophenyl hydrazone (CCCP), a common inducer of mitophagy (i.e., one type of autophagy), was used to damage mitochondria and reduce cell viability. Previous studies have shown that the mitochondrial membrane potential changes rapidly during the CCCP treatment, which results in fluctuations in the fluorescence of a probe^27,45^. Thus, to accurately observe the fluorescence imaging, we incubated CCCP-treated HeLa cells to allow the cells to acclimate to the new microenvironment and thereby make the drug work more efficiently^46-48^. Next, CCCP-treated HeLa cells stained by PCV-1 were imaged on SIM. As shown in **Fig. 6a**, upon the addition of CCCP, the filamentous fluorescence of mitochondria broke and became spherical fluorescence in different sizes, thereby indicating that mitophagy had occurred^46,47^. As the time of CCCP treatment was lengthened, more damaged mitochondria were observed, meaning that the degree of cell viability was decreased. As expected, PCV-1 was located in the nucleolus in dead cells due to the disappearance of mitochondrial membrane potential. PCV-1 escaped from damaged mitochondria into the nucleolus during cell injury and showed bright red fluorescence in both organelles, also as expected. In particular, SIM images after 24 h treatment of CCCP exhibited remarkably high fluorescence intensity in the nucleolus but less spherical fluorescence in the cytoplasm, which further confirms the exceptionally low cell viability under that condition. Moreover, the corresponding distribution maps of 3D fluorescence intensity under the CCCP treatment at different times (**Fig. 6b**) allowed the dynamic observation of PCV-1’s mitochondria-to-nucleolus migration. The significantly high fluorescence intensities of mitochondria in the control sample gradually diminished during the process of cell injury, while a new high-intensity fluorescent peak emerged in the center of the nuclei. Taking all those imaging data together, we anticipate that PCV-1’s migration ability can be used as a marker of cell viability, as depicted in **Fig. 6c**.

**Fig. 6.**
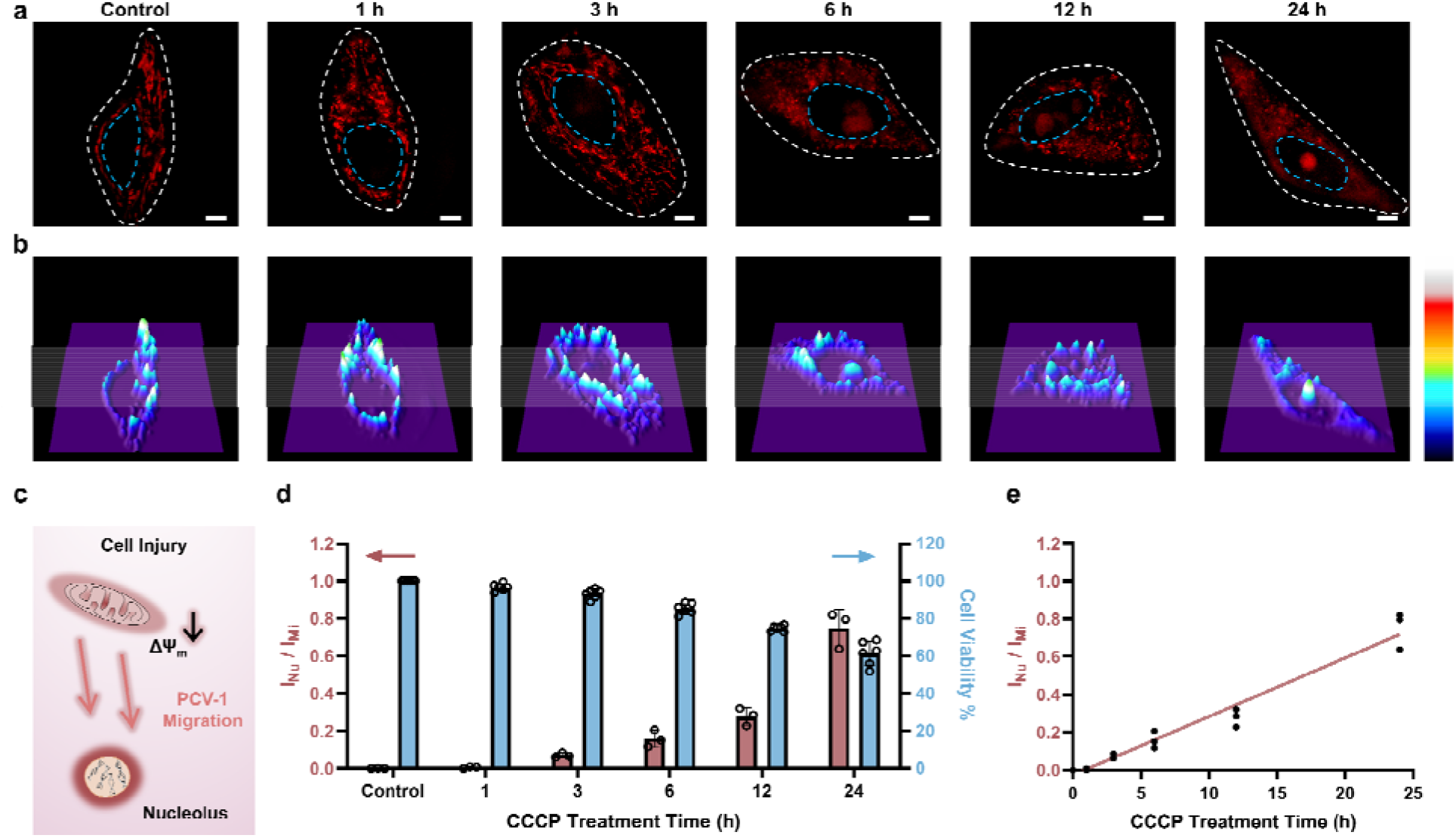
Quantifying cell viability using the ORP analysis with PCV-1. **a** SIM images of HeLa cells stained with PCV-1 after treatment with 20 μM CCCP at different times (0-24 h). PCV-1 channel: Ex 561 nm, Em 570-640 nm; scale bars: 5 μm. **b** 3D fluorescence intensity plots of SIM images in **a**, constructed by ImageJ. **c** Schematic illustration of PCV-1’s migration from mitochondria to nucleolus during cell injury. **d** ORP ratios of I_Nu_/I_Mi_ in PCV-1-stained HeLa cells after treatment with 20 μM CCCP at different times (0-24 h) (red column, data are given a *M ± SEM*, *n* = 3); cell viability measurements of 20 μM CCCP treated HeLa cells at different times (0-24 h) using the conventional CCK-8 assay (blue column). **e** Plots of ORP ratios of I_Nu_/I_Mi_ versus CCCP treatment time, the red line stands for the linear relationship between fluorescence intensity ratio and CCCP treatment time (R^2^ = 0.96). I_Nu_, the fluorescence intensity of nucleolus; I_Mi_, the fluorescence intensity of mitochondria.

As next-generation methods of quantitative cell biology, the digitization of single cells and subcellular organelles has attracted considerable attention in recent years. Nevertheless, systems of quantitative analysis for measuring cell viability have not been developed. Herein, we propose a new method of image analysis, named *organelle ratiometric probing* (ORP), which is defined by calculating fluorescence intensity ratios between various morphologies at the subcellular level, including in mitochondria, lysosomes, and endoplasmic reticulum. Having observed the migration of PCV-1 from mitochondria to the nucleolus, we then used the ORP for quantifying cell viability. Among the ORP’s merits when using PCV-1, long spatial distances between mitochondria and the nucleolus without direct contact allow accurately quantifying the fluorescence intensities of organelles at the single-cell level using super-resolution imaging. Another merit is that single-channel imaging can significantly reduce instrumental errors and permit reasonable comparisons of the fluorescence intensity of organelles in situ. Last, the PCV’s high resistance to photobleaching enables using ORP to observe fluorescence imaging both at multiple times and in the long term without sacrificing any sensitivity.

To digitize SIM images of CCCP-treated HeLa cells, PCV-1’s nucleolus-to-mitochondria fluorescence intensity ratios (i.e., ORP ratio) were calculated and displayed in **Fig. 6d** and **Supplementary Fig. 18**. Moreover, to correlate those fluorescence ratios with cell viability, the cell viability of CCCP-treated HeLa cells was also investigated using the conventional CCK-8 assay, as shown in the right blue columns of **Fig. 6d**. PCV-1 initially exhibited ORP ratios close to 0 owing to the healthy cells with high mitochondrial membrane potential. As treatment with CCCP continued, cell viability gradually decreased, which resulted in an increase in the nucleolus-to-mitochondria fluorescence intensity ratio. Furthermore, as shown in the results of HeLa cells treated with CCCP for 12 h, a fluorescence intensity ratio of 0.3 represents a cell viability of 80%. According to the International Organization for Standardization, a percentage of cell viability less than 80% is considered to indicate cytotoxicity^49^. With that knowledge, we chose 0.3 as the cutoff point for our ORP analysis with PCV-1 to assess whether adding a drug would cause cytotoxicity. In addition, when plotting the fluorescence intensity ratios of nucleolus-to-mitochondria against the duration of CCCP treatment (**Fig. 6e**), the ORP analysis with PCV-1 demonstrated an exceptional linear relationship (*R*^2^ = 0.96), providing additional evidence that it is an excellent ratiometric tool for evaluating cell viability. Next, the HeLa cells stained by PCV-1 but without the treatment of CCCP were studied as well. As shown in **Supplementary Fig. 19a**, no migration from mitochondria to the nucleolus was observed, and PCV-1 exhibited stable, strong filamentous fluorescence after 24 h of incubation. Moreover, as shown in **Supplementary Fig. 10**, PCV-1 did not affect cell viability after 24 h of incubation. Therefore, per the ORP analysis, the PCV-1 fluorescence ratios for the SIM images were consistently less than 0.3, which further demonstrates that PCV-1 is non-cytotoxic (**Supplementary Fig. 19b**, **20**).

### Quantifying cell viability under different concentrations of CCCP treatment

Next, to confirm that PCV-1 is indeed an excellent tool for dynamically visualizing cell viability, we also investigated CCCP concentration-dependent autophagy. As shown in **Fig. 7a**, as the concentration of CCCP increased, the number of damaged mitochondria increased as well, and more PCV-1 accumulated in the nucleolus, thereby suggesting that autophagy occurred more intensely under pathological conditions^46,47^. However, if the concentration of CCCP was too high, then the strong cytotoxicity would destroy the cells, such that PCV-1 displayed only a significantly high fluorescence peak in the nucleolus (**Fig. 7b**). These results imply that PCV-1 can vividly display the states of cells at different stages in the process of cell injury. Interpreting those SIM images using the ORP analysis with PCV-1 (**Supplementary Fig. 21**) revealed nucleolus-to-mitochondria intensity ratios indicating a growing trend with the increase of the CCCP concentration, as illustrated in **Fig. 7c**. For the range of each fluorescence ratio, there was a corresponding range of cell viability to match. Thus, when cell viability was less than 80% in the experiment with the treatment of 10 μM CCCP, PCV-1’s fluorescence ratio exceeded 0.3, which further confirmed that our assumed cutoff point of 0.3 was appropriate. Additionally, the fluorescence intensity ratios measured by the ORP analysis with PCV-1 also exhibited a decent linear correlation as the concentration of CCCP treatment increased (**Fig. 7d**). In sum, in super-resolution imaging, PCV-1 together with the ORP analysis allowed visualizing cell injury and quantifying cell viability at the single-cell level.

**Fig. 7.**
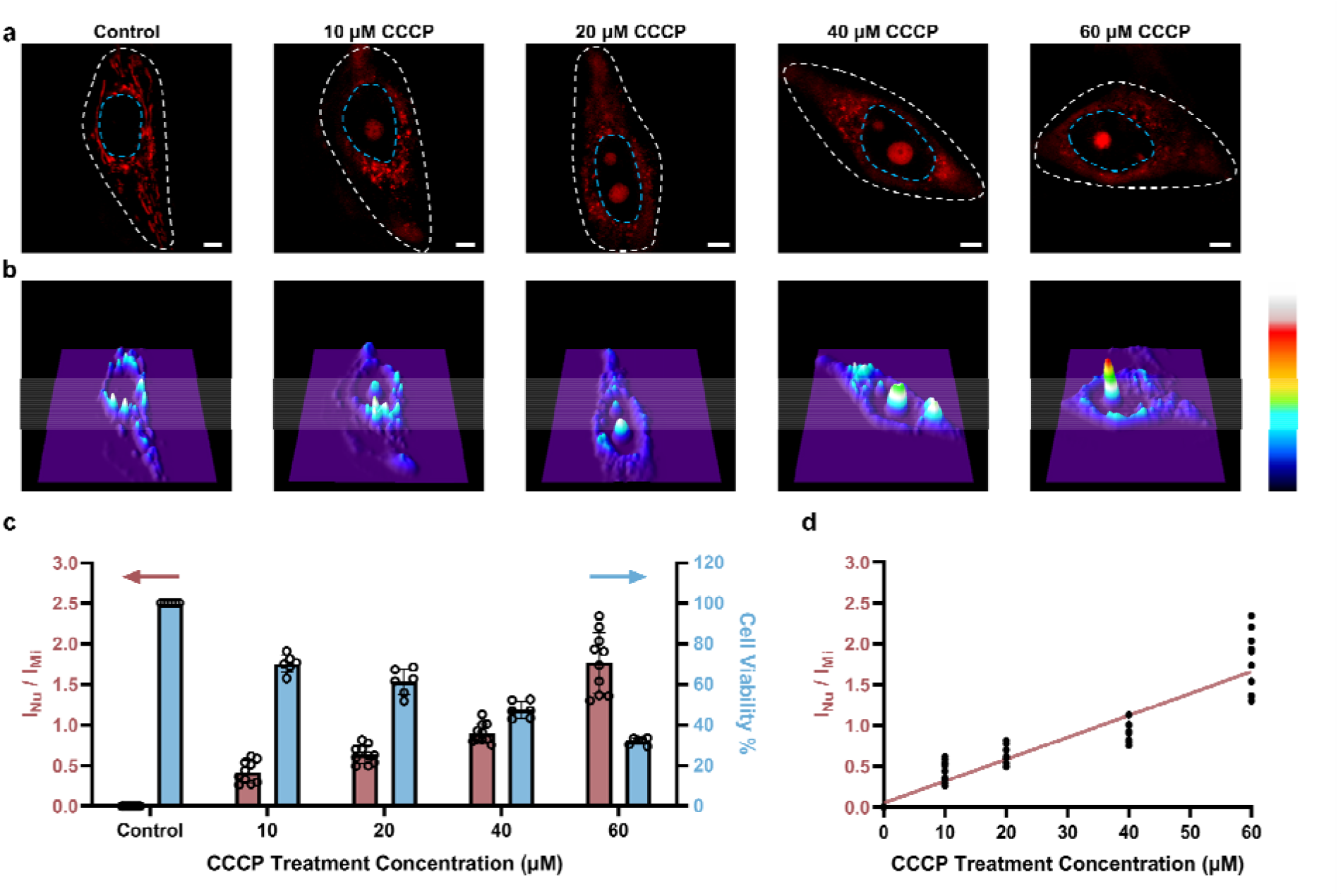
Quantifying cell viability under different concentrations of CCCP treatment. **a** SIM images of HeLa cells stained with PCV-1 after treatment with different concentrations of CCCP (0-60 μM) for 24 h. PCV-1 channel: Ex 561 nm, Em 570-640 nm; scale bars: 5 μm. **b** 3D fluorescence intensity plots of SIM images in **a**, constructed by ImageJ. **c** ORP ratios of I_Nu_/I_Mi_ in PCV-1-stained HeLa cells after treatment with different concentrations of CCCP (0-60 μM) for 24 h (red column, data are given as *M ± SEM*, *n* = 10); cell viability measurements of different concentrations of CCCP (0-60 μM) treated HeLa cells for 24 h using the commercial CCK-8 assay (blue column). **d** Plots of ORP ratios of I_Nu_/I_Mi_ versus CCCP treatment concentration, the red line stands for the linear relationship between fluorescence intensity ratio and CCCP treatment concentration (R^2^ = 0.87).

### Visualizing cell viability in various routes toward cell death using the ORP analysis

To explore the applications of the ORP analysis in different routes toward cell death, we investigated the SIM images of PCV-1-stained HeLa cells after various drug treatments. One route was hydrogen peroxide (H_2_O_2_), a reactive oxygen species that can efficiently inhibit metabolism and cause cell damage by inducing either apoptosis or necrosis depending on the dose^50,51^. Another route was sodium azide (NaN_3_), a highly toxic salt that acts as a potent inhibitor of mitochondrial respiration by hindering the function of cytochrome oxidase and induces cell death^52-54^. The last one was chloroquine, an inhibitor of the autophagy–lysosome pathway that has been tested extensively in preclinical cancer models and used as an antimalarial drug^55,56^. In all, H_2_O_2_, NaN_3_, and chloroquine were applied to treat HeLa cells to induce cell death.

As shown in **Fig. 8a**, the control group without drug treatment presented clearly filamentous fluorescence of PCV-1 in mitochondria. As the high concentration of H_2_O_2_ treatment induced cell necrosis, PCV-1 quickly lit up the nucleolus, whereas the high fluorescence in the cytoplasm reduced rapidly due to the low cell viability under this condition (**Fig. 8b**). Meanwhile, the NaN_3_-treated HeLa cells demonstrated spherical fluorescence in both the cytoplasm and nucleolus, thereby suggesting that cell death via apoptosis can be visualized by PCV-1 (**Fig. 8c**). PCV-1 also allows monitoring the low viability of chloroquine-treated cells, as reflected in intense fluorescence in the center of nuclei (**Fig. 8d**). To probe cell viability after drug treatments in greater depth, we additionally checked those SIM images in the ORP analysis. All HeLa cells treated with drugs exhibited fluorescence intensity ratios of the nucleolus to the mitochondria exceeding 1.5, as shown in **Fig. 8e**. With such high fluorescence intensity ratios, we can confidently propose that H_2_O_2_, NaN_3_, and chloroquine are highly cytotoxic, a condition that extremely reduces cell viability.

**Fig. 8.**
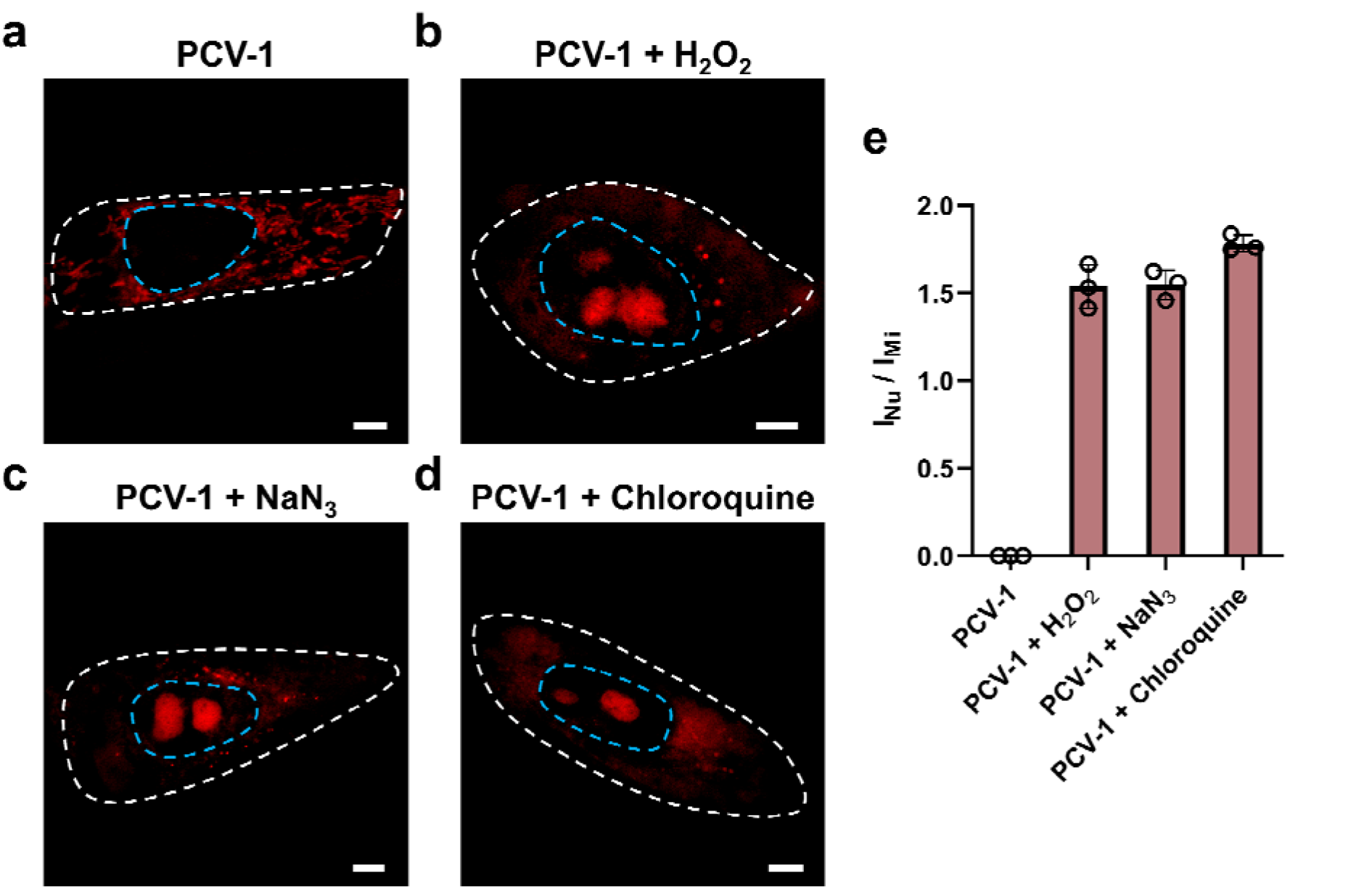
Visualizing cell viability in various routes toward cell death using the ORP analysis. **a** SIM images of HeLa cells stained with PCV-1. SIM images of HeLa cells stained with PCV-1 after treatment of **b** 50 mM H_2_O_2_ for 30 min, **c** 0.5% NaN_3_ for 20 min, **d** 50 μM Chloroquine for 30 min. Scale bars: 5 μm. **e** ORP ratios of I_Nu_/I_Mi_ in PCV-1-stained HeLa cells after treatment with different drugs. Data are given as *M ± SEM*, *n* = 3.

Last, we explored SIM images of PCV-1 in other cell lines during CCCP-induced mitophagy. As shown in **Supplementary Fig. 22**, PCV-1 migrated from mitochondria to the nucleolus in both HDFn and MCF-7 cells, thereby indicating that it can be adapted to other cell lines to visualize cell viability. In that light, PCV-1 can serve as a versatile tool for rapidly quantifying cell viability in different cells and various pathways toward cell death, including apoptosis, autophagy, and necrosis.

In this study, we rationally designed and synthesized a fluorescent probe, PCV-1, to visualize cell viability under SIM. Because of PCV-1’s outstanding photophysical properties and excellent super-resolution imaging performance, its migration from mitochondria to the nucleolus could be dynamically visualized at the single-cell level during cell injury. By extension, harnessing PCV-1’s excellent photostability and signal-to-noise ratio and by comparing the fluorescence intensity of the two organelles, mitochondria and nucleoli, we developed a powerful analytical assay, ORP, that we applied to quantitatively analyze and efficiently assess the viability of individual cells, thereby enabling deeper insights into the potential mechanisms of cell death. In our ORP analysis with PCV-1, we identified 0.3 as the cutoff point for assessing whether adding the drug will cause apparent cytotoxicity, which greatly expands the probe’s applicability. In the future, combined with machine learning and big data analysis, our proposed ORP analysis with PCV-1 stands to significantly improve the diagnosis of disease, facilitate the development of drugs, and reduce medical costs.

## Methods

### Materials

All commercially available chemicals were purchased and used without further treatment, including methanol, ethanol, glycerol, acetone, acetonitrile, chloroform, dimethylformamide (DMF), dimethyl sulfoxide (DMSO), hexane, 6-bromo-2-methylquinoline, phenylboronic acid, methyl iodide, *p*-fluorobenzaldehyde, 1-methylpiperazine. LysoTracker Deep Red (LTDR, #L12492), MitoTracker Red FM (MTR, #M22425), MitoTracker Deep Red FM (MTDR, #M22426), Hoechst 33342 (#H3570), and DAPI were purchased from Invitrogen (Thermo Fisher Scientific). All fluorescent dyes were used according to the product manuals. Carbonyl cyanide *m*-chlorophenylhydrazone (CCCP, #C2759) was purchased from Sigma. Cell Counting Kit-8 (CCK-8) assay was purchased from Dojindo Molecular Technologies. Penicillin, streptomycin, fetal bovine serum (FBS), and Dulbecco’s modified Eagle’s medium (DMEM) were all purchased from Gibco (Thermo Fisher Scientific). Phosphate-buffered saline (PBS) was purchased from Hyclone (GE Healthcare Life Sciences). Calf thymus DNA solution was purchased from ThermoFisher.

### Characterization

The ^1^H and ^13^C NMR spectra were recorded on a Bruker Avance III HD Ascend 400 MHz NMR spectrometer. Chemical shifts for protons are referenced to the residual solvent peak (CDCl_3_, ^1^H NMR: 7.26 ppm; D_2_O, ^1^H NMR: 4.79 ppm; *d_6_*-DMSO, ^1^H NMR: 2.50 ppm), while chemical shifts for carbons are referenced to the residual solvent peaks (CDCl_3_, ^13^C NMR: 77.16 ppm, *d_6_*-DMSO; ^13^C NMR: 39.51 ppm). The following abbreviations (or combinations thereof) were used to explain multiplicities: s = singlet, d = doublet, m = multiplet. Matrix-Assisted Laser Desorption Ionization (MALDI) mass spectrometry was performed on a Bruker Biflex III MALDI-TOFMS instrument.

## Synthesis

### Synthesis of 2-methyl-6-phenylquinoline

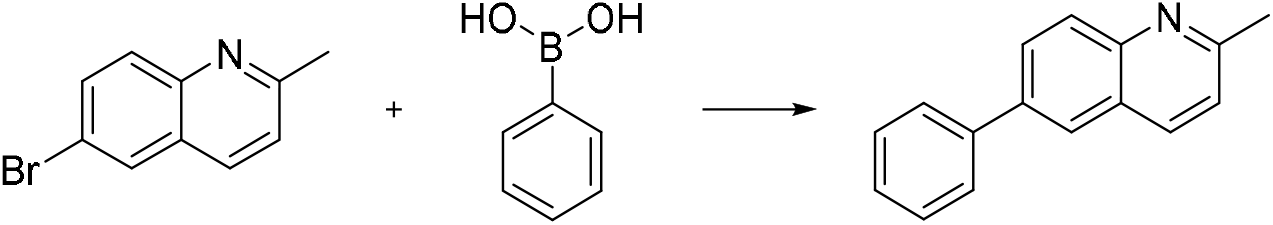

To a three-neck round flask, 5.0 mmol (1.11 g) 6-bromo-2-methylquinoline, 1.5 equiv. phenylboronic acid, 0.1 equiv. Pd(PPh_3_)_4_, 3.0 equiv. K_2_CO_3_, and 0.1 equiv. Bu_4_NBr were added under N_2_, then 180 mL toluene, 120 mL EtOH, and 60 mL H_2_O were added. The reaction mixture was stirred at 100 °C overnight. When the reaction was finished, the mixture was diluted with EtOAc and washed with water. The combined organic phase was concentrated. The pure product was obtained through chromatography on silica using EtOAc, EtOH, and hexanes as the eluent. Eventually, 0.72 g target product was obtained with a yield of 65%. ^1^H NMR (400 MHz, CDCl_3_) δ 8.11-8.08 (m, 2 H), 7.96-7.94 (m, 2 H), 7.73-7.70 (m, 2 H), 7.51-7.47 (m, 2 H), 7.41-7.37 (m, 1 H), 7.32 (d, 1 H), 2.77 (s, 3 H). ^13^C NMR (100 MHz, CDCl_3_) δ 159.16, 147.36, 145.59, 138.58, 136.54, 129.28, 129.15, 129.06, 127.70, 127.51, 126.79, 125.39, 122.56, 25.51.

### Synthesis of 1,2-dimethyl-6-phenylquinolin-1-ium

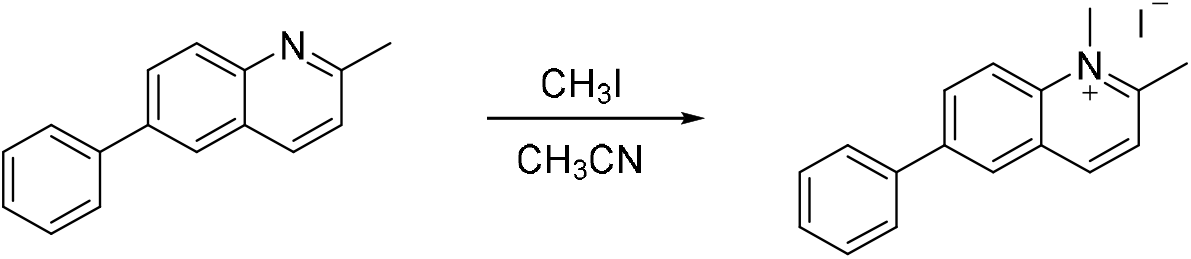

To a 25 mL round flask, 0.5 mmol (110 mg) 2-methyl-6-phenylquinoline and 2.0 equiv. CH_3_I were dissolved in 10 mL dry CH_3_CN. The reaction mixture was stirred at 80 °C overnight. After being filtered and washed with cool ethanol and dimethyl ether, the pure product was obtained through recrystallization from ethanol. Finally, 40 mg desired product was obtained (yield = 23%). ^1^H NMR (400 MHz, DMSO) δ 9.10 (d, 1 H), 8.73 (d, 1 H), 8.66 (d, 1 H), 8.59-8.56 (m, 1 H), 8.13 (d, 1 H), 7.96-7.94 (m, 2 H), 7.60 (m, 2 H), 7.52 (m, 1 H), 4.47 (s, 3 H), 3.08 (s, 3 H). ^13^C NMR (100 MHz, DMSO) δ 160.86, 145.48, 140.08, 138.63, 137.20, 133.67, 129.41, 129.03,

128.37, 127.35, 127.13, 125.55, 119.79. 22.99.

### Synthesis of 4-(4-methylpiperazin-1-yl) benzaldehyde

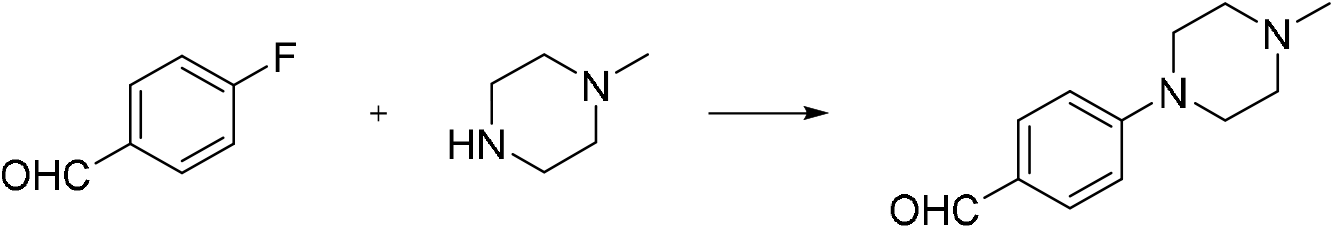

10 mmol *p*-fluorobenzaldehyde (1.24 g), 1.1 equiv. 1-methylpiperazine (1.10 g), 2.0 equiv. K_2_CO_3_ (2.76 g), and DMF (100 mL) were added to a round flask. The mixture was stirred at 100 °C for 16 h under N_2_. After the reaction was completed, the mixture was poured into cool water and precipitation was formed. The precipitation was filtered and washed with cool water. After drying at room temperature, the pure product was obtained (1.98 g, yield of 97%) through chromatography on silica with EtOAc and hexanes as the eluent. ^1^H NMR (400 MHz, CDCl_3_) δ 9.76 (s, 1 H), 7.74 (d, 2 H), 6.90 (d, 2 H), 3.40 (t, 4 H), 2.54 (t, 4 H), 2.34 (s, 3 H). ^13^C NMR (100 MHz, CDCl_3_) δ 190.52, 155.11, 131.95, 127.21, 113.64, 54.78, 47.16, 46.22.

### Synthesis of 1-methyl-2-(4-(4-methylpiperazin-1-yl)styryl)-6-phenylquinolin-1-ium (PCV-1)

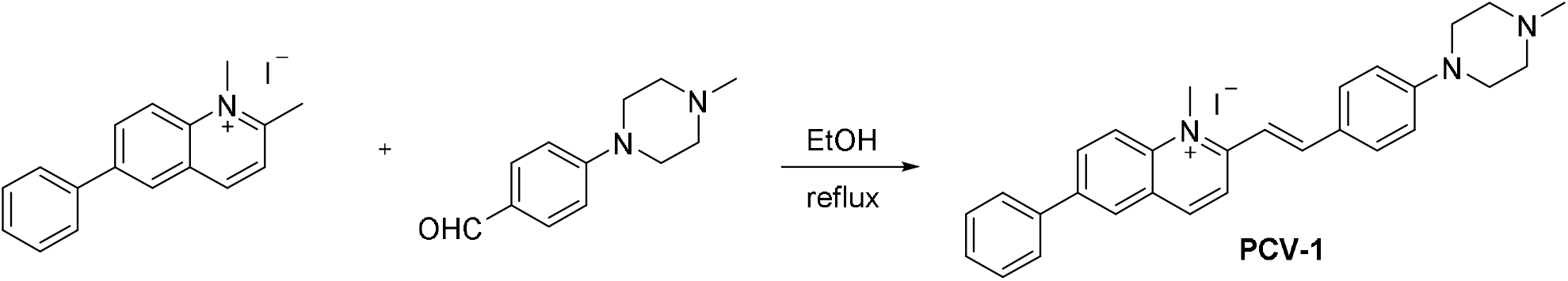

To a round bottom flask, 0.1 mmol 1,2-dimethyl-6-phenylquinolin-1-ium (35.0 mg) and 1.1 equiv. 4-(4-methylpiperazin-1-yl) benzaldehyde (22.4 mg) were dissolved in 10 mL anhydrous ethanol. The solution was refluxed for 48 h under N_2_. After cooling to room temperature, the solvent was evaporated under reduced pressure. The residue was purified by washing with cool ethanol, EtOAc, hexanes, and dimethyl ether. Finally, 25.0 mg desired product was obtained with a yield of 45%. ^1^H NMR (400 MHz, DMSO) δ 8.62 (d, 1 H), 8.50 (d, 1 H), 8.19 (s, 2 H), 8.10 (m, 1 H), 8.00 (d, 1 H), 7.85 (d, 2 H), 7.70 (m, 2 H), 7.61 (d, 1 H), 7.52 (m, 2 H), 7.45 (m, 1 H), 4.65 (s, 3 H), 3.35 (m, 4 H), 2.55 (m, 4 H), 2.37 (s, 3 H). ^13^C NMR (100 MHz, DMSO) δ 148.73, 142.43, 138.63, 137.46, 134.86, 133.84, 132.56, 129.44, 127.57, 127.23, 126.86, 124.68, 122.08, 119.24, 115.45, 113.93, 112.93, 106.55, 94.27, 54.34, 46.62, 41.52, 29.71.

### Absorption and emission measurements

PCV-1 in aqueous solution (10 μM) was transferred to a quartz cuvette with 1 cm optical length to measure the absorption spectra on an Agilent Cary 8453 spectrophotometer. The fluorescence spectra were recorded on a HORIBA Fluorolog QM spectrofluorometer.

### The measurements of lipophilicity

Lipophilicity was determined by a shake-flask ultraviolet spectrophotometry method to measure the *n*-octanol/water partition coefficients (logP).^57^ First, 25 mL 1-octanol and 25 mL aqueous buffer solution were mixed at room temperature. After 24 h, the standard solutions of the sample (10 μM) were prepared by using the preceding saturated 1-octanol phase and aqueous buffer phase solution, respectively. Next, the test solutions were prepared by mixing 10 ml standard solution in 1-octanol and 10 ml standard solution in aqueous buffer solution. After shaking, the mixture of test solution was centrifuged for 5 min. The 1-octanol phase layer was separated from the water phase layer and the absorbance of the two layers was measured on an Agilent Cary 8453 spectrophotometer. Based on the Beer-Lambert law, the concentrations of the sample in the 1-octanol phase (*C_o_*) and the aqueous phase (*C_w_*) were calculated. Finally, the n-octanol/water partition coefficients (logP) were further calculated by using the following equation. logP = log (*C_o_/C_w_*)

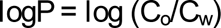

### F□rster–Hoffmann equation

The linear relationship between log *I_605 nm_* and log η. F□rster– Hoffmann equation: log *I* = *x* log η + C, where I is the emission intensity, x is the sensitivity of the probe towards viscosity, η is the viscosity, and C is a constant.^29,38^

### Photostability test

The photostability test of probes was investigated under a light irradiation wavelength of 488 nm for 300 s.

### Phosphate buffer solution

A series of standard pH buffer solutions were prepared by mixing 0.2 M KH_2_PO_4,_ 0.2 M Na_2_HPO_4_·7H_2_O, and 0.2 M H_3_PO_4_ with different volume ratios, and the pH values were measured using a Mettler Toledo FiveEasy Plus™ pH-meter.

### Computational methods of DFT

All calculations were performed with the Gaussian 16W program package employing the DFT method with Becke’s three-parameter hybrid functional and Lee-Yang-Parr’s gradient corrected correlation functional (B3LYP).^58-60^ 6-31G* basis set was applied for H, C, and N.^61^ The geometries of the singlet ground states of compounds were optimized in H_2_O using the conductive polarizable continuum model (CPCM). The local minimum on each potential energy surface was confirmed by frequency analysis. Time-dependent DFT calculations produced the singlet excited states of each compound starting from the optimized geometry of the corresponding singlet ground state, using the CPCM method with H_2_O as the solvent. The calculated absorption spectra, electronic transition contributions, and electron density difference maps (EDDMs) were generated by GaussSum 3.0.^62^ The electronic orbitals were visualized using VMD 1.9.4. Isovalue was 0.04 for plotting HOMOs, LUMOs, and EDDMs.^63^

### Molecular docking calculations

The molecular docking calculations were performed using the AutoDock Vina software. ^44^ The structure of PCV-1 was obtained from previous DFT calculations via Gaussian 16W software. The initial structures of DNA were downloaded from the protein data bank (PDB ID: 1D30).^64^

### Cell culture

HeLa cells were gifted from Dr. Carolyn M. Price lab (University of Cincinnati). Cells were cultured in DMEM containing 10% FBS, 100 units mL^-1^ streptomycin, and 100 units mL^-1^ penicillin. The cells were cultured in the incubator (Thero Fisher Scientific) at 37°C with 5% CO_2_ and 100% humidity.

### Structured illumination microscopy (SIM)

Super-resolution images were performed on a commercial Nikon Structured Illumination Microscopy (N-SIM, version AR5.11.00 64 bit, Tokyo, Japan) super-resolution miscroscope. Images were obtained at 512 × 512 by using the Nikon NIS-Elements software, and the raw images were reconstructed and analyzed with Nikon NIS-Elements and ImageJ. Cells were cultured on a 35 × 10 mm dish for 24 h for imaging. Cells were seeded on culture dishes for 24 h to adhere. The colocalization experiments were obtained with a dual-channel imaging mode. HeLa cells were stained by PCV-1 (0.4 μM, 30 min). The Pearson’s colocalization coefficient (PCC) values were analyzed and calculated using the CellProfiler software. Quantification of fluorescence intensity ratios was collected via ImageJ and Excel.

### Cell viability and cytotoxicity assay (CCK-8)

HeLa cells were seeded in a 96-well plate with a density of 1 × 10^4^ cells/well in 100 µL culture medium. After 24 h to adhere, different concentrations of PCV-1 were added to the well plate and incubated for 24 h. Next, 10 µL CCK-8 was added to each cell well plate for incubating for 1 h. Next, the absorbance at 490 nm was obtained using a microplate reader (BioTek Instruments, Inc., USA). The viability of CCCP-treat cells were also measured with the same method.

## Acknowledgments

Y.S. acknowledges the support of the National Science Foundation (CHE1955358) and University of Cincinnati. J.D. is supported by the National Institute of Health (R35GM128837).

## Author Contributions

R.C. characterized the photophysical properties, performed bioimaging, and analyzed all the imaging data. K.Q. helped process the imaging data. G.H. performed the DFT calculations. B.K. performed molecular docking calculations. G.D. synthesized probe. Y.S. and J.D. conceived the project, designed the experiments, and wrote the manuscript with the help of all authors.

## References

1. Hotchkiss, R. S., Strasser, A., McDunn, J. E. & Swanson, P. E. Cell death. New Engl. J. Med. 361, 1570–1583, (2009).

2. Elmore, S. Apoptosis: a review of programmed cell death. Toxicol. Pathol. 35, 495–516, (2007).

3. Evan, G. I. & Vousden, K. H. Proliferation, cell cycle and apoptosis in cancer. Nature 411, 342–348, (2001).

4. Kepp, O., Galluzzi, L., Lipinski, M., Yuan, J. & Kroemer, G. Cell death assays for drug discovery. Nat. Rev. Drug Discov. 10, 221–237, (2011).

5. Tian, M., Ma, Y. & Lin, W. Fluorescent probes for the visualization of cell viability. Acc. Chem. Res. 52, 2147–2157, (2019).

6. Banfalvi, G. Methods to detect apoptotic cell death. Apoptosis 22, 306–323, (2017).

7. Muppidi, J., Porter, M. & Siegel, R. M. Measurement of apoptosis and other forms of cell death. Curr. Protoc. Immunol. 59, 3.17.11–13.17.36, (2004).

8. Berridge, M. V., Herst, P. M. & Tan, A. S. in Biotechnol. Annu. Rev. Vol. 11 127–152 (Elsevier, 2005).

9. Larsson, P. et al. Optimization of cell viability assays to improve replicability and reproducibility of cancer drug sensitivity screens. Sci. Rep. 10, 5798, (2020).

10. Zorova, L. D. et al. Mitochondrial membrane potential. Anal. Biochem. 552, 50–59, (2018).

11. Green, D. R. & Reed, J. C. Mitochondria and apoptosis. Science 281, 1309–1312, (1998).

12. Tian, M., Sun, J., Dong, B. & Lin, W. Dynamically monitoring cell viability in a dual-color mode: construction of an aggregation/monomer-based probe capable of reversible mitochondria-nucleus migration. Angew. Chem. Int. Ed. 57, 16506-16510, (2018).

13. Li, X. et al. Spatially dependent fluorescent probe for detecting different situations of mitochondrial membrane potential conveniently and efficiently. Anal. Chem. 89, 3335–3344, (2017).

14. Fang, H. et al. Super-resolution imaging of mitochondrial HClO during cell ferroptosis using a near-infrared fluorescent probe. Anal. Chem. 94, 17904–17912, (2022).

15. Wang, L., et al. Super-resolution analyzing spatial organization of lysosomes with an organic fluorescent probe. Exploration, (2022).

16. Qiu, K. et al. Light-activated mitochondrial fission through optogenetic control of mitochondria-lysosome contacts. Nat. Commun. 13, 4303, (2022).

17. Fang, H. et al. Simultaneous Zn^2+^ tracking in multiple organelles using super-resolution morphology-correlated organelle identification in living cells. Nat. Commun. 12, 109, (2021).

18. Chen, Q. et al. A dual-labeling probe to track functional mitochondria–lysosome interactions in live cells. Nat. Commun. 11, 6290, (2020).

19. Satori, C. P. et al. Bioanalysis of eukaryotic organelles. Chem. Rev. 113, 2733–2811, (2013).

20. Wang, K.-N. et al. A continuous add-on probe reveals the nonlinear enlargement of mitochondria in light-activated oncosis. Adv. Sci. 8, 2004566, (2021).

21. Fang, H. et al. De novo-designed near-infrared nanoaggregates for super-resolution monitoring of lysosomes in cells, in whole organoids, and in vivo. ACS Nano 13, 14426–14436, (2019).

22. Qiu, K., et al. Ultralong-term super-resolution tracking of lysosomes in brain organoids by near-infrared noble metal nanoclusters. ACS Mater. Lett., 1565-1573, (2022).

23. Qiu, K. et al. De novo design of a membrane-anchored probe for multidimensional quantification of endocytic dynamics. Adv. Healthc. Mater. 11, 2102185, (2022).

24. Yang, R., et al. A single fluorescent pH probe for simultaneous two-color visualization of nuclei and mitochondria and monitoring cell apoptosis. ACS Sens. 6, 1552-1559, (2021)

25. Tian, M., Zhan, J. & Lin, W. Single fluorescent probes enabling simultaneous visualization of duple organelles: design principles, mechanisms, and applications. Coord. Chem. Rev. 451, (2022).

26. Wang, K. N., et al. A polarity-sensitive ratiometric fluorescence probe for monitoring changes in lipid droplets and nucleus during ferroptosis. Angew. Chem. Int. Ed. Engl. 60, 15095-15100, (2021).

27. Tian, M., Sun, J., Dong, B. & Lin, W. Construction of mitochondria-nucleolus shuttling fluorescent probe for the reversible detection of mitochondrial membrane potential. Sensors Actuators B: Chem. 292, 16–23, (2019).

28. Zou, Z. et al. Real-time visualizing mitophagy-specific viscosity dynamic by mitochondria-anchored molecular rotor. Anal. Chem. 91, 8574–8581, (2019).

29. Chen, B. et al. Sensing and imaging of mitochondrial viscosity in living cells using a red fluorescent probe with a long lifetime. Chem. Commun. 55, 7410–7413, (2019).

30. Abeywickrama, C. S. Large stokes shift benzothiazolium cyanine dyes with improved intramolecular charge transfer (ICT) for cell imaging applications. Chem. Commun. 58, 9855–9869, (2022).

31. Mastro, A. M., Babich, M. A., Taylor, W. D. & Keith, A. D. Diffusion of a small molecule in the cytoplasm of mammalian cells. Proc. Natl. Acad. Sci. U.S.A. 81, 3414–3418, (1984).

32. Dijksterhuis, J., Nijsse, J., Hoekstra, F. A. & Golovina, E. A. High viscosity and anisotropy characterize the cytoplasm of fungal dormant stress-resistant spores. Eukaryot. Cell 6, 157–170, (2007).

33. Luby-Phelps, K., Taylor, D. L. & Lanni, F. Probing the structure of cytoplasm. J. Cell Biol. 102, 2015–2022, (1986).

34. Wang, L., Xiao, Y., Tian, W. & Deng, L. Activatable rotor for quantifying lysosomal viscosity in living cells. J. Am. Chem. Soc. 135, 2903–2906, (2013).

35. Yang, Z. et al. A self-calibrating bipartite viscosity sensor for mitochondria. J. Am. Chem. Soc. 135, 9181–9185, (2013).

36. Li, S.-J. et al. A dual-response fluorescent probe for the detection of viscosity and H_2_S and its application in studying their cross-talk influence in mitochondria. Anal. Chem. 90, 9418–9425, (2018).

37. Zhu, H. et al. A “distorted-bodipy”-based fluorescent probe for imaging of cellular viscosity in live cells. Chem. Eur. J. 20, 4691–4696, (2014).

38. Song, Y. et al. One stone, three birds: pH triggered transformation of aminopyronine and iminopyronine based lysosome targeting viscosity probe for cancer visualization. Anal. Chem. 93, 1786–1791, (2021).

39. Bastiaens, P. I. H. & Squire, A. Fluorescence lifetime imaging microscopy: spatial resolution of biochemical processes in the cell. Trends Cell Biol. 9, 48–52, (1999).

40. Horobin, R. W. & Rashid-Doubell, F. Predicting small molecule fluorescent probe localization in living cells using qsar modeling. 2. Specifying probe, protocol and cell factors; selecting qsar models; predicting entry and localization. Biotech. Histochem. 88, 461-476, (2013).

41. Horobin, R. W., Rashid-Doubell, F., Pediani, J. D. & Milligan, G. Predicting small molecule fluorescent probe localization in living cells using qsar modeling. 1. Overview and models for probes of structure, properties and function in single cells. Biotech. Histochem. 88, 440-460, (2013).

42. Qiu, K. et al. Biscylometalated iridium(III) complexes target mitochondria or lysosomes by regulating the lipophilicity of the main ligands. Dalton Trans. 45, 16144–16147, (2016).

43. Mukherjee, T. et al. Live-cell imaging of the nucleolus and mapping mitochondrial viscosity with a dual function fluorescent probe. Org. Biomol. Chem. 19, 3389–3395, (2021).

44. Trott, O. & Olson, A. J. AutoDock Vina: improving the speed and accuracy of docking with a new scoring function, efficient optimization, and multithreading. J. Comput. Chem. 31, 455–461, (2010).

45. Li, M.-Y. et al. A mitochondria–nucleolus migration fluorescent probe for monitoring of mitochondrial membrane potential and identification of cell apoptosis. Anal. Methods 11, 5750–5754, (2019).

46. Chen, Q. et al. Nanoscale monitoring of mitochondria and lysosome interactions for drug screening and discovery. Nano Res. 12, 1009–1015, (2019).

47. Shao, X. et al. Super-resolution quantification of nanoscale damage to mitochondria in live cells. Nano Res. 13, 2149–2155, (2020).

48. Qiu, K. et al. Super-resolution observation of lysosomal dynamics with fluorescent gold nanoparticles. Theranostics 10, 6072–6081, (2020).

49. López-García, J., Lehocký, M., Humpolíček, P. & Sáha, P. Hacat keratinocytes response on antimicrobial atelocollagen substrates: extent of cytotoxicity, cell viability and proliferation. J. Funct. Biomater. 5, 43–57 (2014).

50. Saito, Y. et al. Turning point in apoptosis/necrosis induced by hydrogen peroxide. Free Radic. Res. 40, 619–630, (2006).

51. Tochigi, M. et al. Hydrogen peroxide induces cell death in human trail-resistant melanoma through intracellular superoxide generation. Int. J. Oncol. 42, 863–872, (2013).

52. Inomata, K. & Tanaka, H. Protective effect of benidipine against sodium azide-induced cell death in cultured neonatal rat cardiac myocytes. J. Pharmacol. Sci. 93, 163–170, (2003).

53. Chen, S. J., Bradley, M. E. & Lee, T. C. Chemical hypoxia triggers apoptosis of cultured neonatal rat cardiac myocytes: modulation by calcium-regulated proteases and protein kinases. Mol. Cell. Biochem. 178, 141–149, (1998).

54. Bowler, M. W., Montgomery, M. G., Leslie, A. G. W. & Walker, J. E. How azide inhibits ATP hydrolysis by the F-ATPases. Proc. Natl. Acad. Sci. U.S.A. 103, 8646–8649, (2006).

55. Fang, H., Liu, A., Dahmen, U. & Dirsch, O. Dual role of chloroquine in liver ischemia reperfusion injury: reduction of liver damage in early phase, but aggravation in late phase. Cell Death Dis. 4, e694–e694, (2013).

56. Gallagher, L. E. et al. Lysosomotropism depends on glucose: a chloroquine resistance mechanism. Cell Death Dis. 8, e3014–e3014, (2017).

57. Qiu, K., Chen, Y., Rees, T. W., Ji, L. & Chao, H. Organelle-targeting metal complexes: from molecular design to bio-applications. Coord. Chem. Rev. 378, 66–86, (2019).

58. Lee, C., Yang, W. & Parr, R. G. Development of the colle-salvetti correlation-energy formula into a functional of the electron density. Phys. Rev. B 37, 785–789, (1988).

59. Becke, A. D. Density-functional thermochemistry. I. The effect of the exchange-only gradient correction. J. Chem. Phys. 96, 2155–2160, (1992).

60. Becke, A. D. Density-functional exchange-energy approximation with correct asymptotic behavior. Phys. Rev. A 38, 3098–3100, (1988).

61. Hehre, W. J. Ab initio molecular orbital theory. Acc. Chem. Res. 9, 399–406, (1976).

62. O’Boyle, N. M., Tenderholt, A. L. & Langner, K. M. Cclib: A library for package-independent computational chemistry algorithms. J. Comput. Chem. 29, 839–845, (2008).

63. Humphrey, W., Dalke, A. & Schulten, K. VMD: visual molecular dynamics. J. Mol. Graph. 14, 33–38, (1996).

64. Larsen, T. A., Goodsell, D. S., Cascio, D., Grzeskowiak, K. & Dickerson, R. E. The structure of DAPI bound to DNA. J. Biomol. Struct. Dyn. 7, 477–491, (1989).

