## Supporting Figures for "Quantifying cell viability through organelle ratiometric probing"

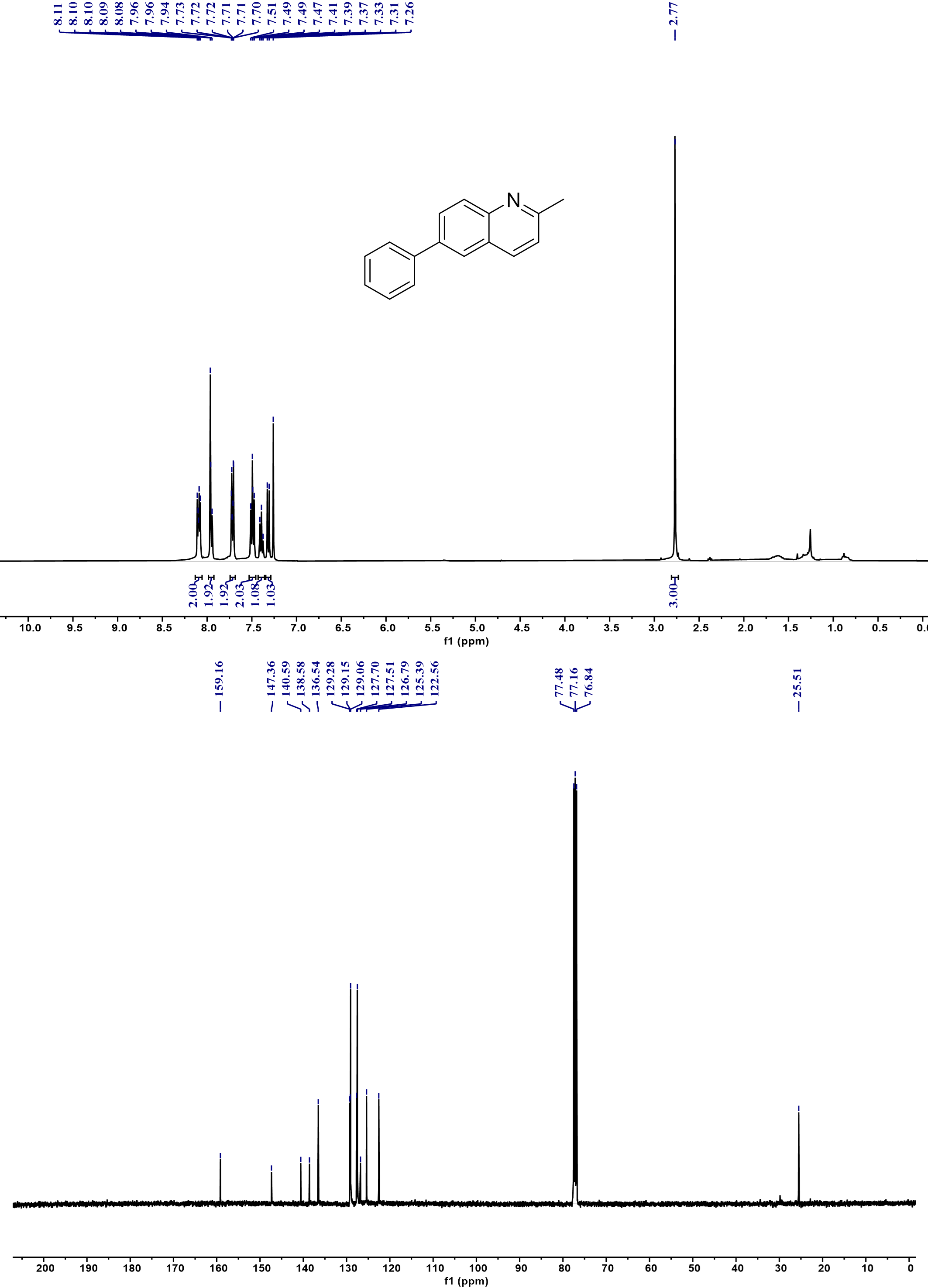

**Supplementary Fig. 1** ^1^H NMR (top) and ^13^C NMR (below) spectra of 2-methyl-6-phenylquinoline.

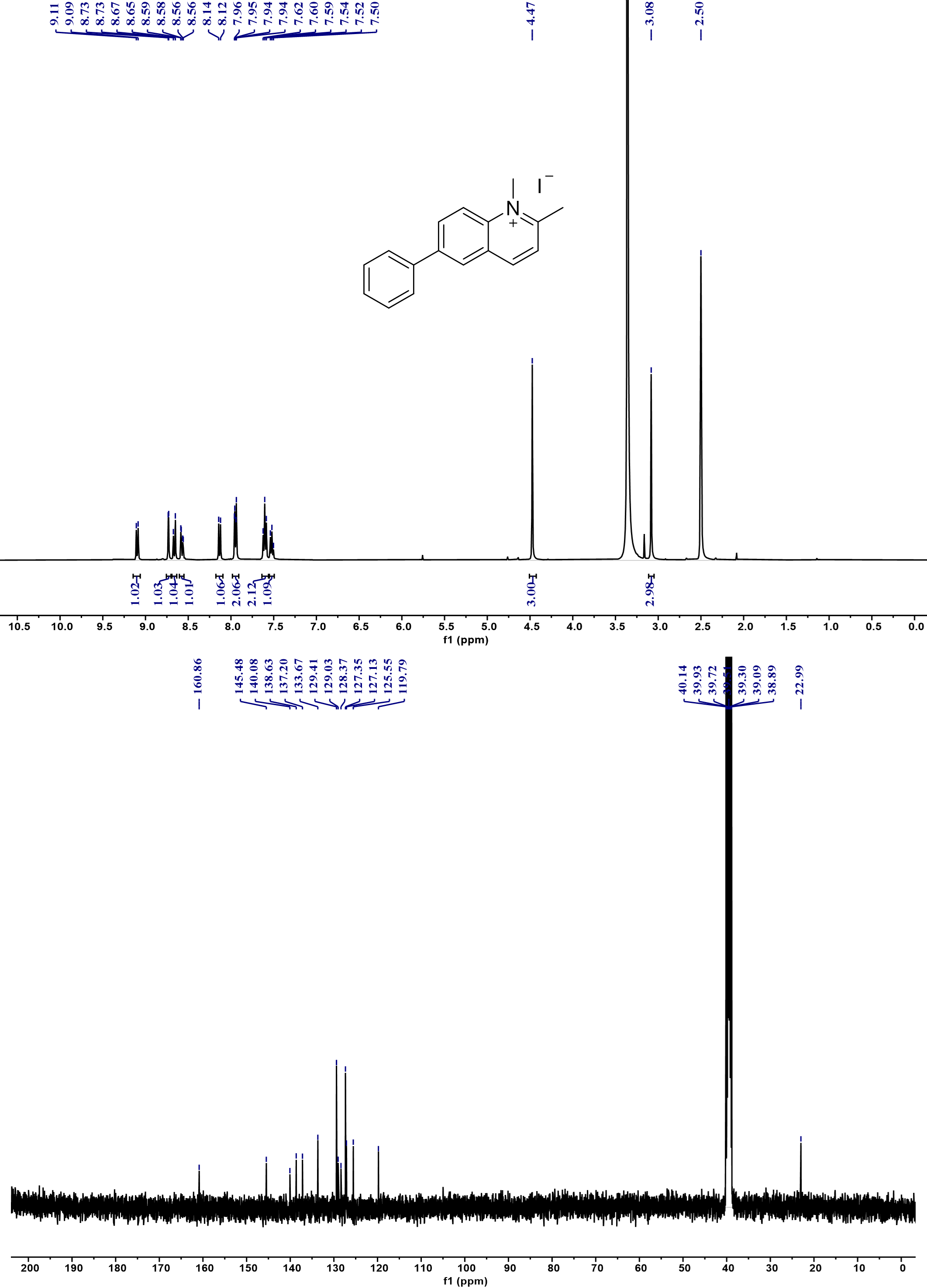

**Supplementary Fig. 2** ^1^H NMR (top) and ^13^C NMR (below) spectra of 1,2-dimethyl-6-phenylquinolin-1-ium.

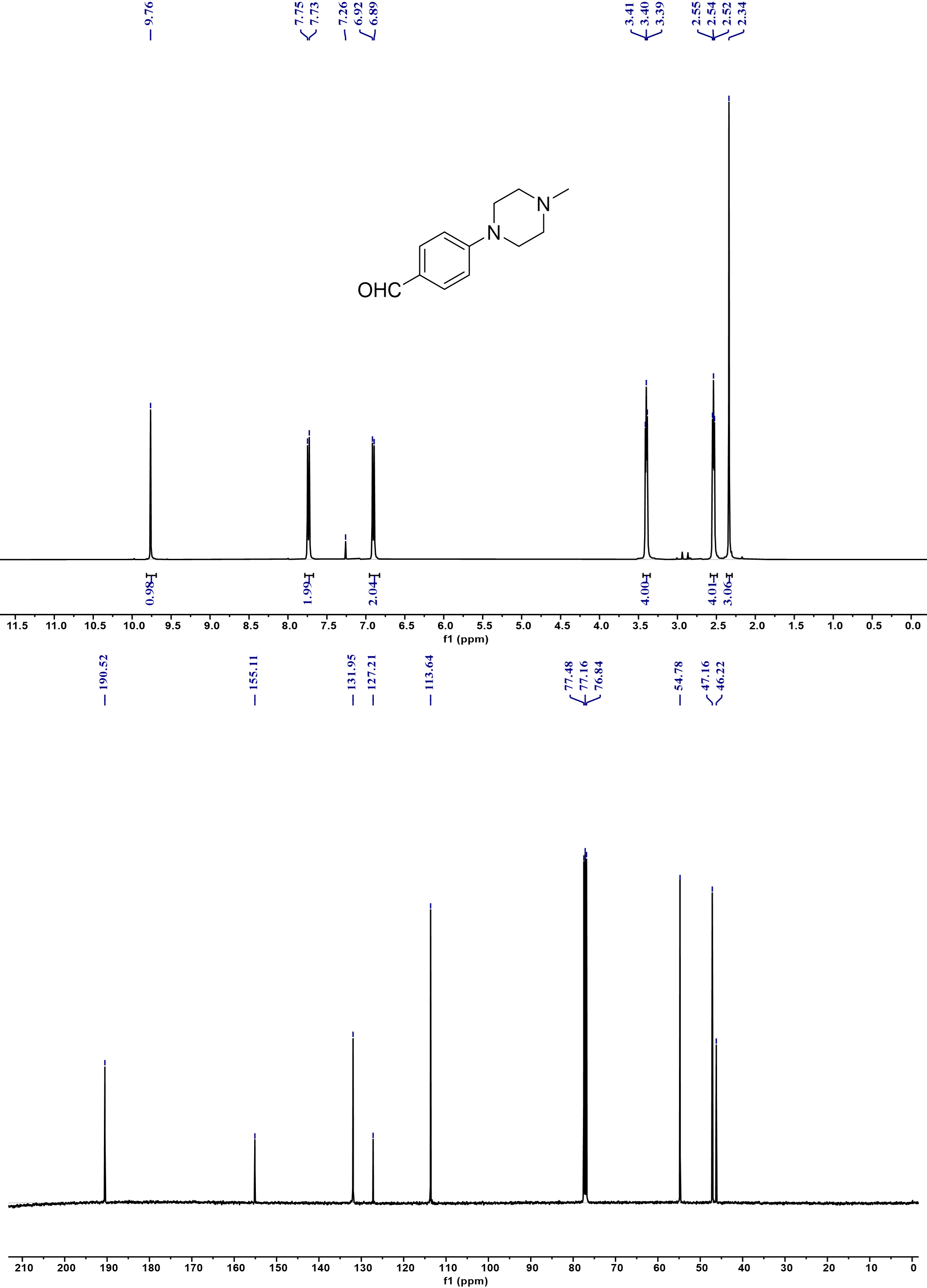

**Supplementary Fig. 3** ^1^H NMR (top) and ^13^C NMR (below) spectra of 4-(4-methylpiperazin-1-yl) benzaldehyde.

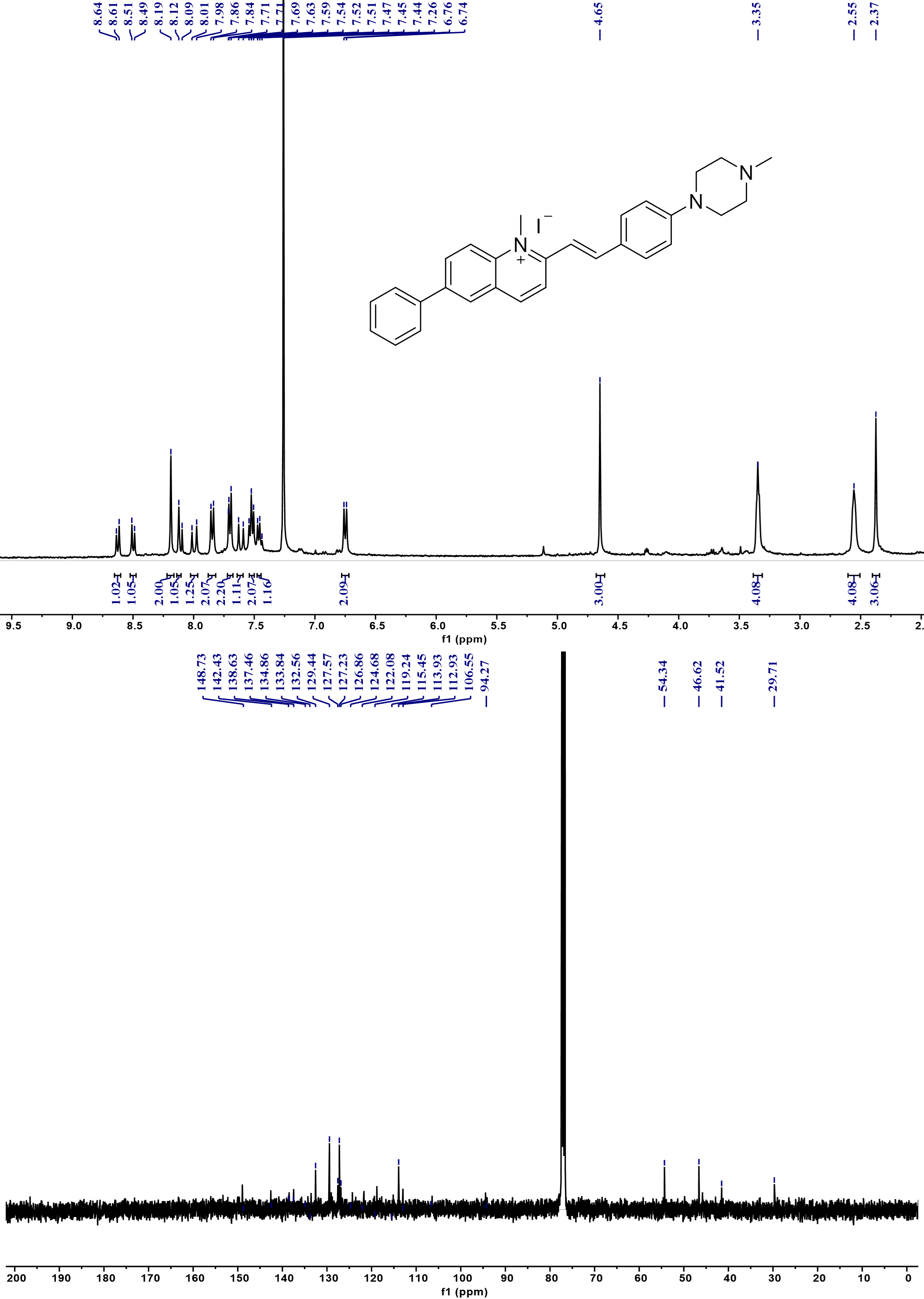

**Supplementary Fig. 4** ^1^H NMR (top) and ^13^C NMR (below) spectra of PCV-1.

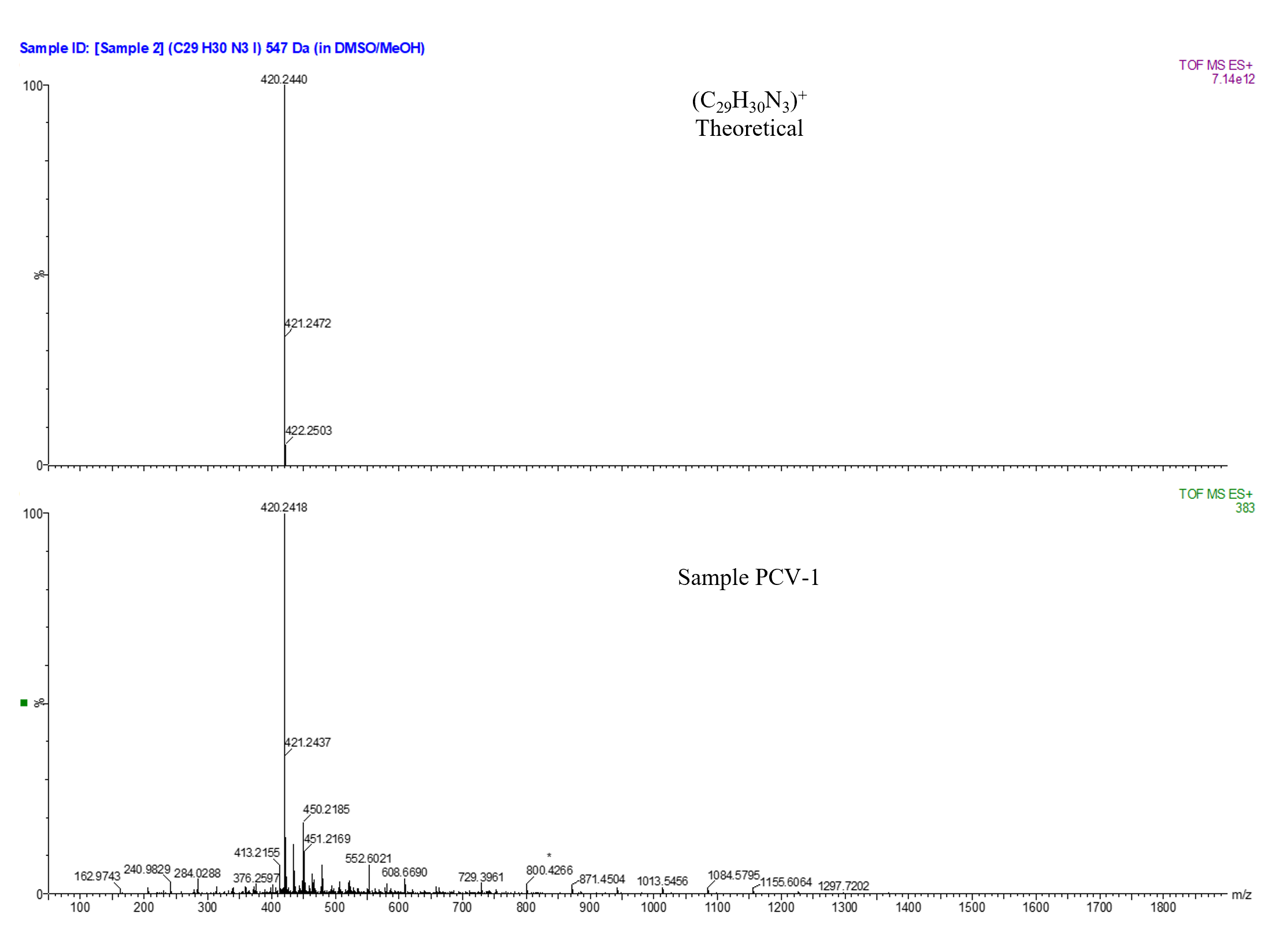

**Supplementary Fig. 5** Mass spectra of PCV-1.

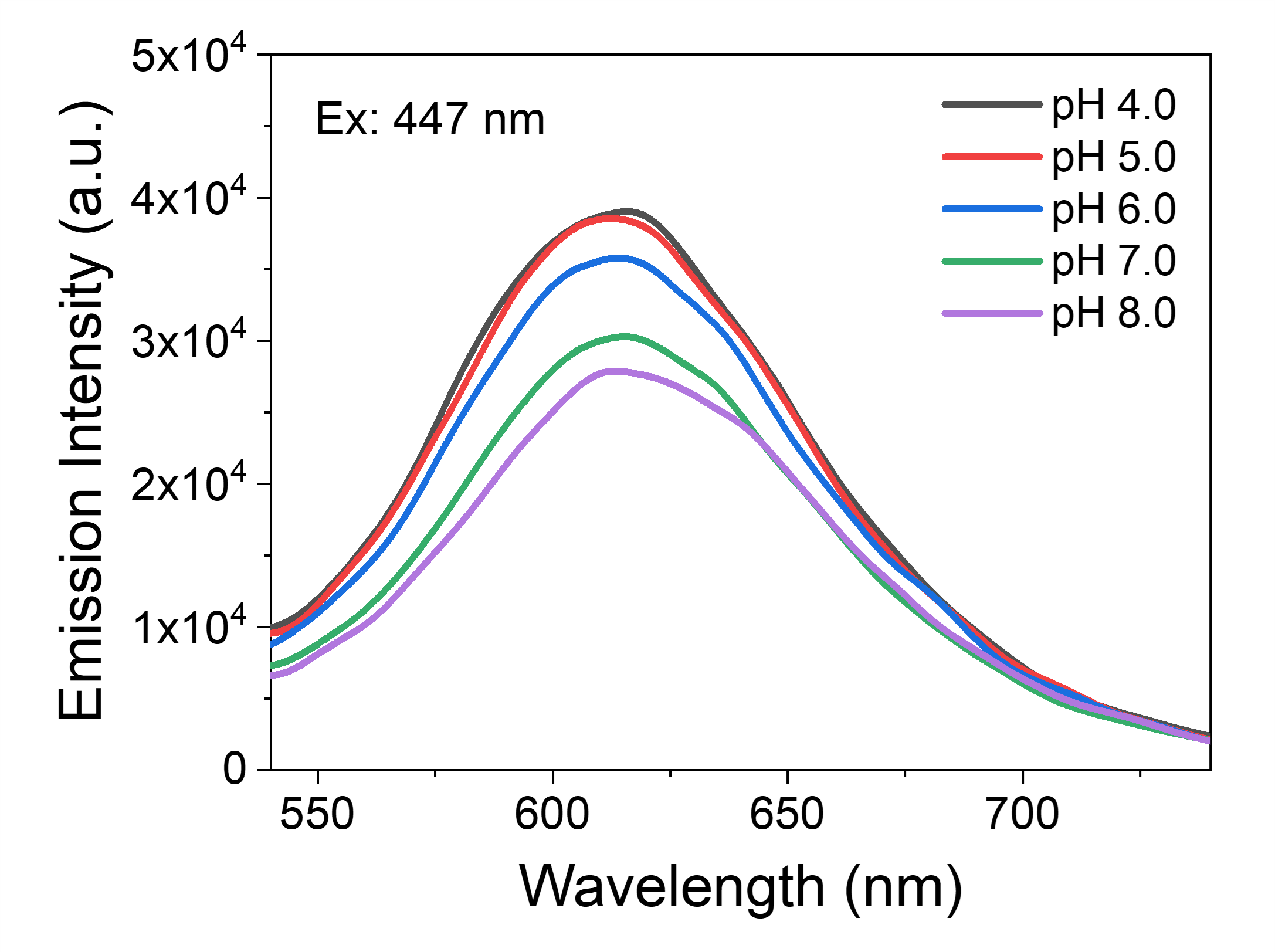

**Supplementary Fig. 6** Emission spectra of PCV-1 in buffer solutions with different pH values.

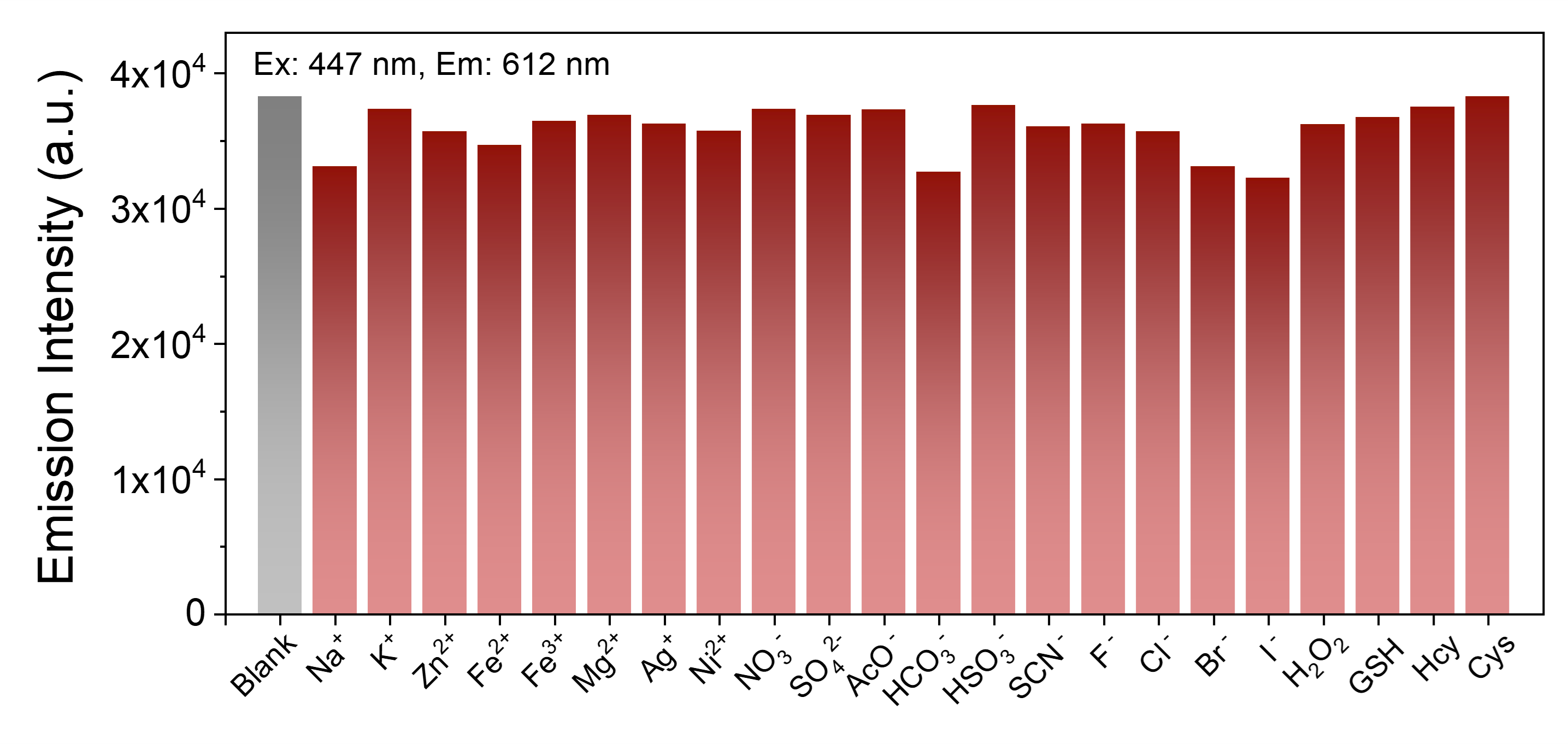
 **Supplementary Fig. 7** Emission intensity histogram plots of PCV-1 in aqueous solution with additional chemical species. Conditions: the concentration of Na^+^ to H_2_O_2_ is 100 μM, and the concentration of GSH to Cys is 200 μM.

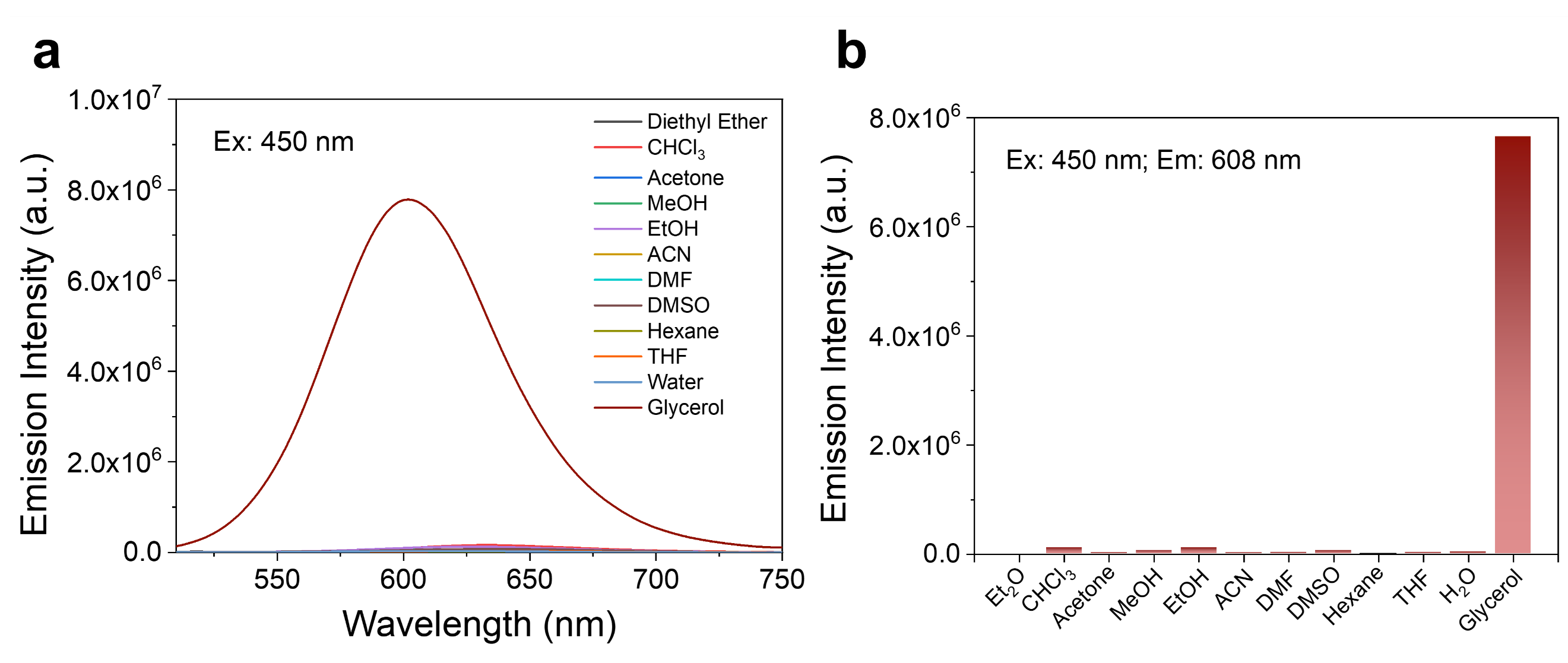

**Supplementary Fig. 8** **a** Emission spectra of PCV-1 in different solvents. **b** Emission intensity histogram plots of PCV-1 in different solvents.

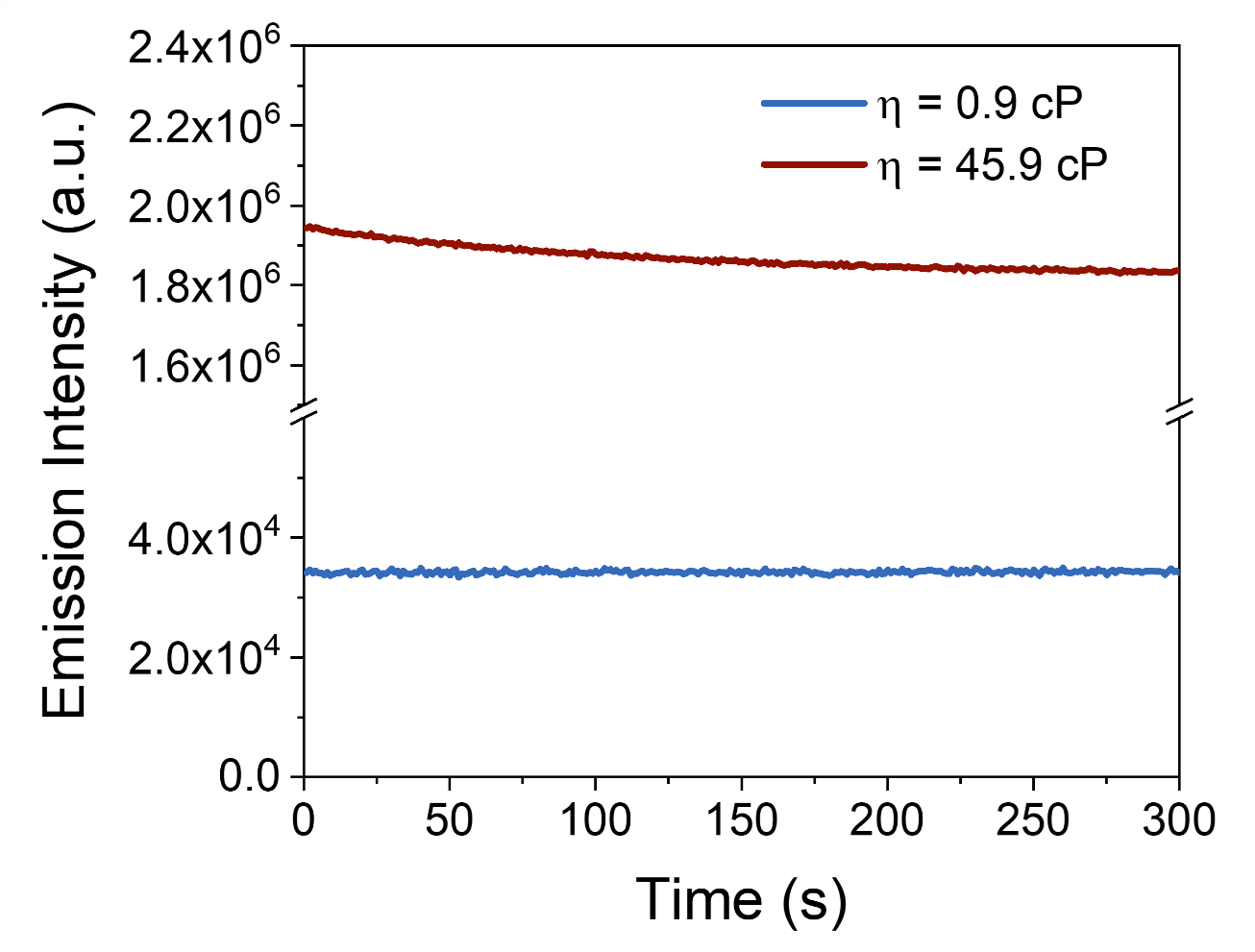

**Supplementary Fig. 9** The photostability tests of PCV-1 in aqueous and high viscosity (49 cP) solutions, *λ_ex_* = 488 nm, *λ_em_* = 610 nm.

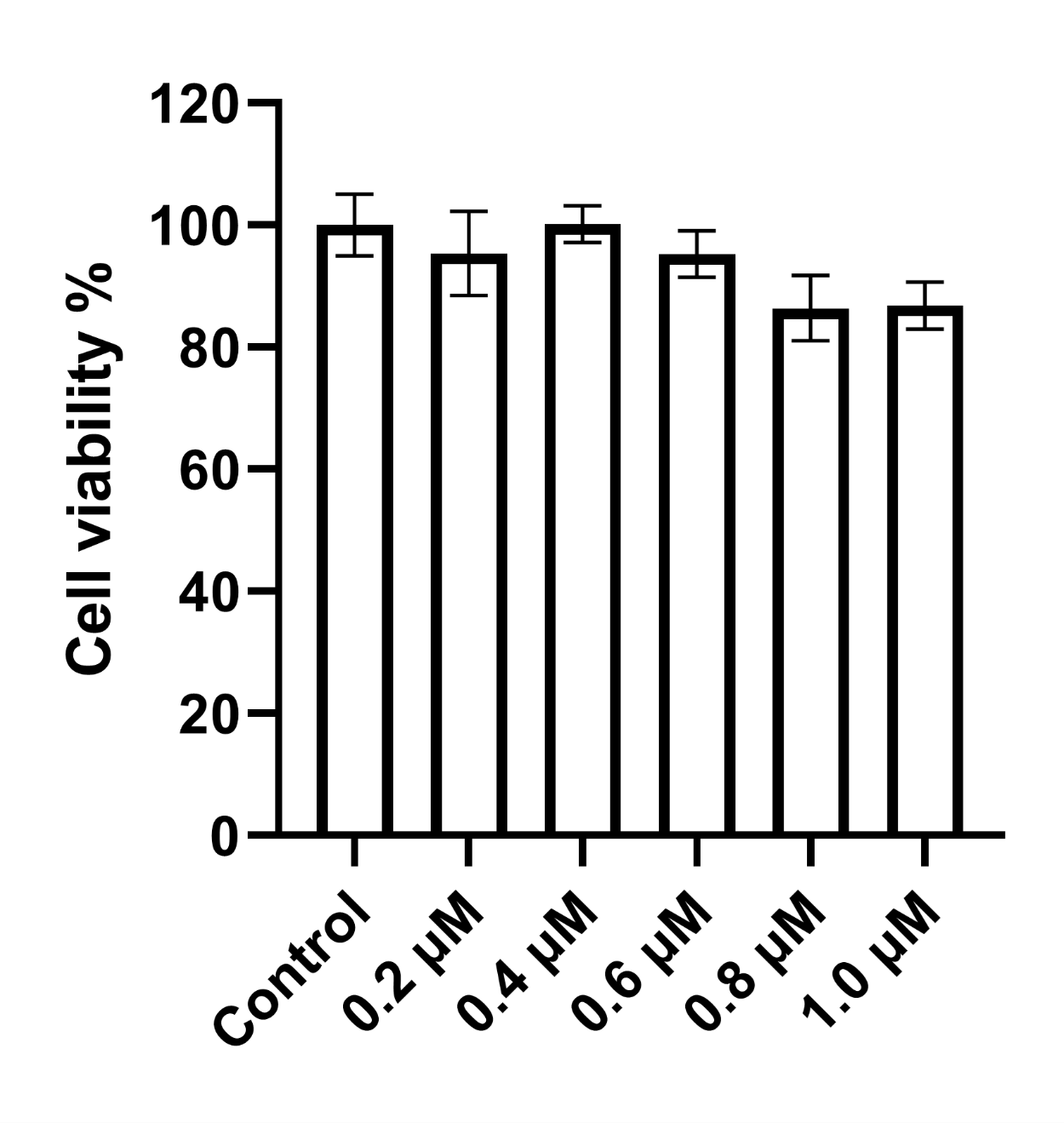

**Supplementary Fig. 10** Cell viability test after HeLa cells were treated with different concentrations of PCV-1 for 24 hours.

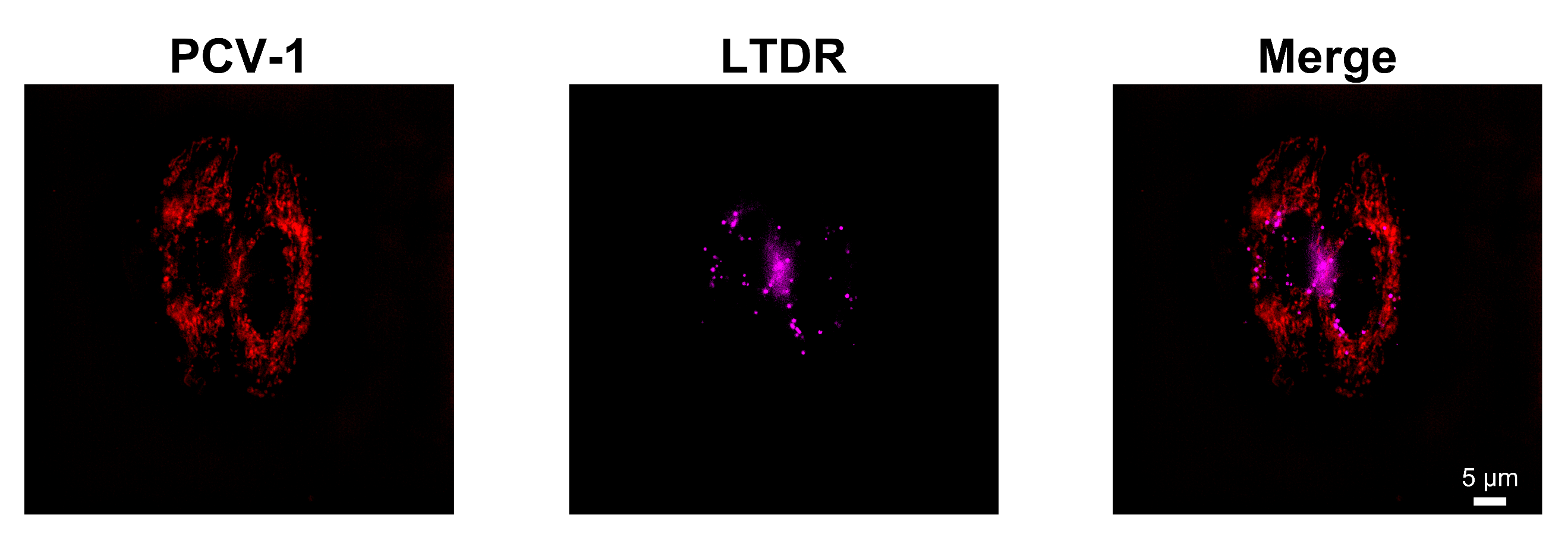

**Supplementary Fig. 11** SIM images of HeLa cells co-stained with PCV-1 and LTDR.

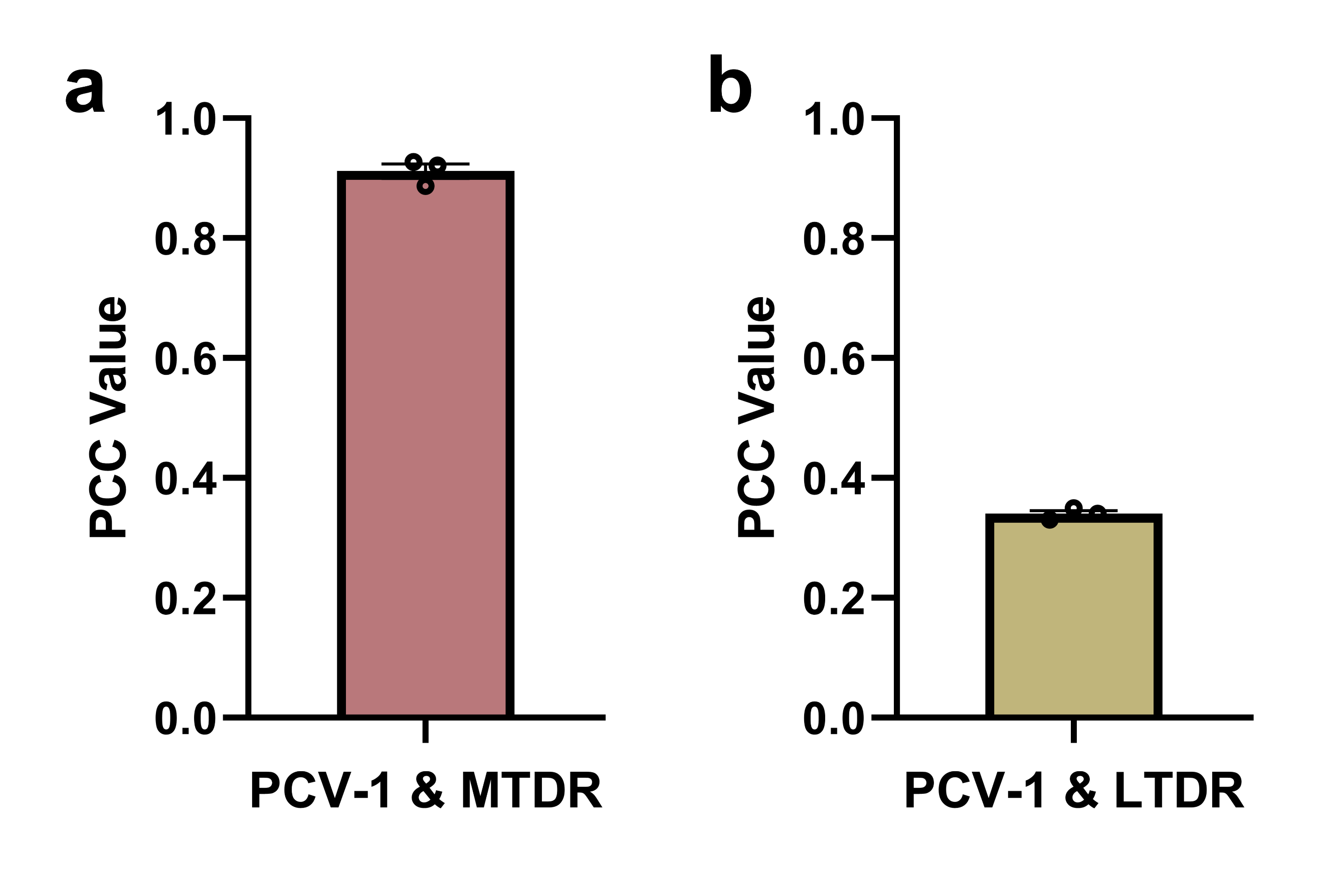

**Supplementary Fig. 12** **a** PCC values of PCV-1 with MTDR. **b** PCC values of PVC-1 with LTDR. Data are given as *M ± SEM*, *n* = 3.

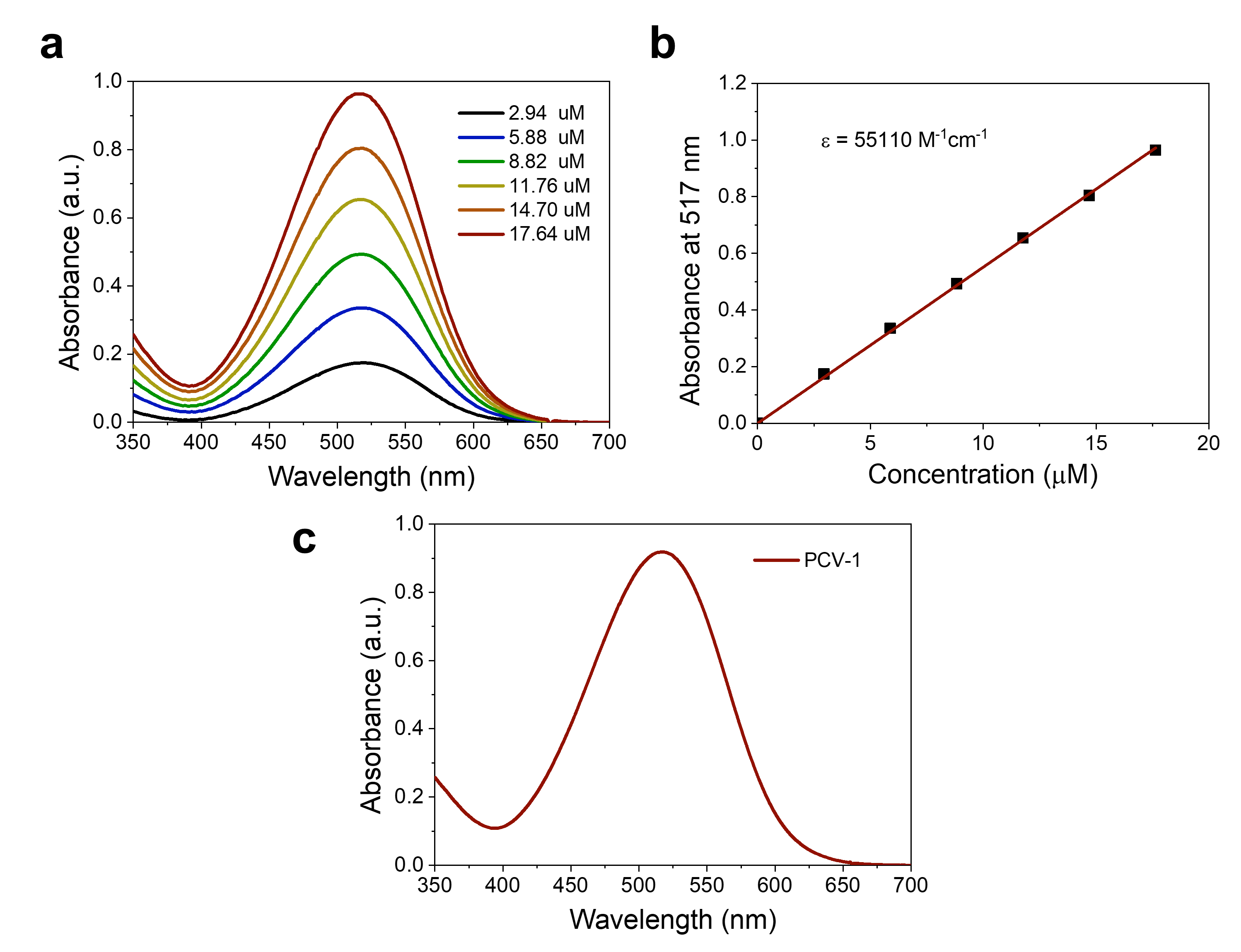

**Supplementary Fig. 13** The measurement of lipophilicity (partition coefficient). **a** Absorption spectra of different concentrations of PCV-1 in PBS buffer solution containing saturated 1-octanol. **b** Scatter plots and linear relationship between absorbance at 517 nm and concentrations of PCV-1. **c** Absorption spectrum of PCV-1 in PBS buffer solution containing saturated 1-octanol after phase separation.

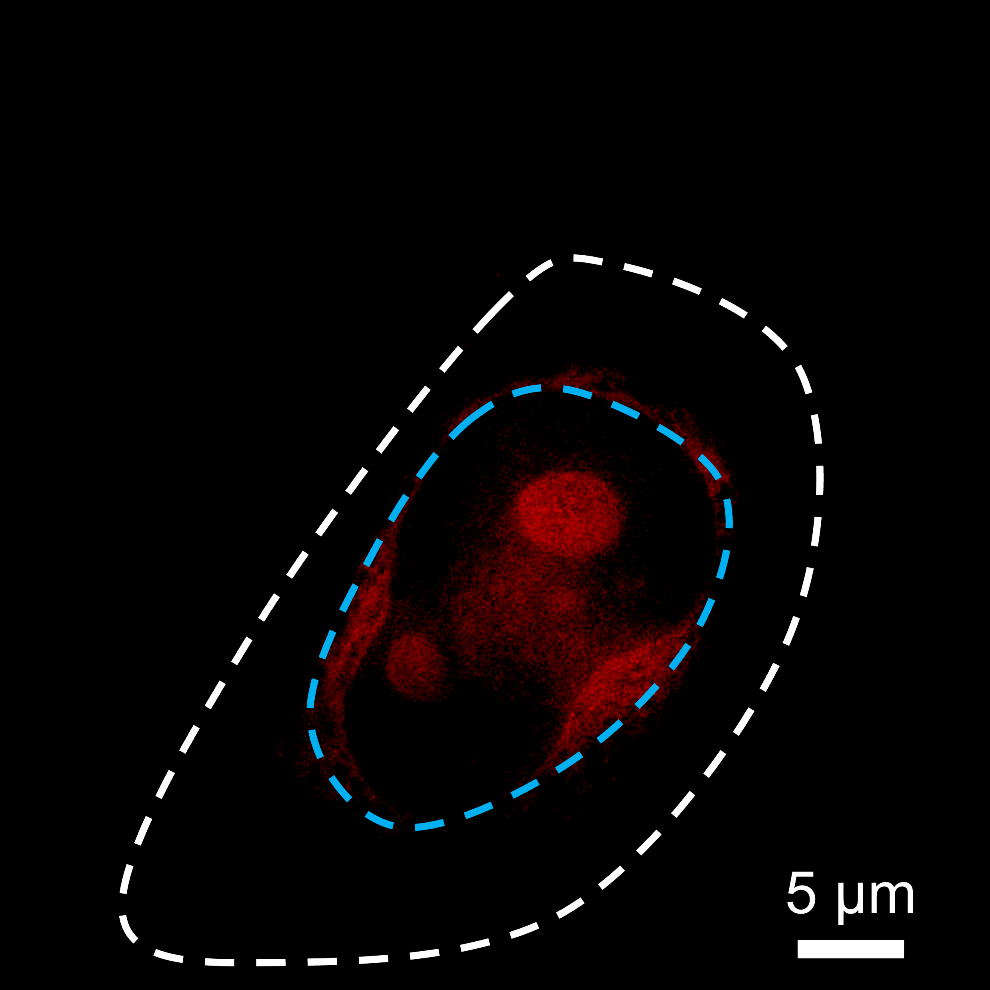

**Supplementary Fig. 14** SIM images of dead HeLa cells stained with PCV-1.

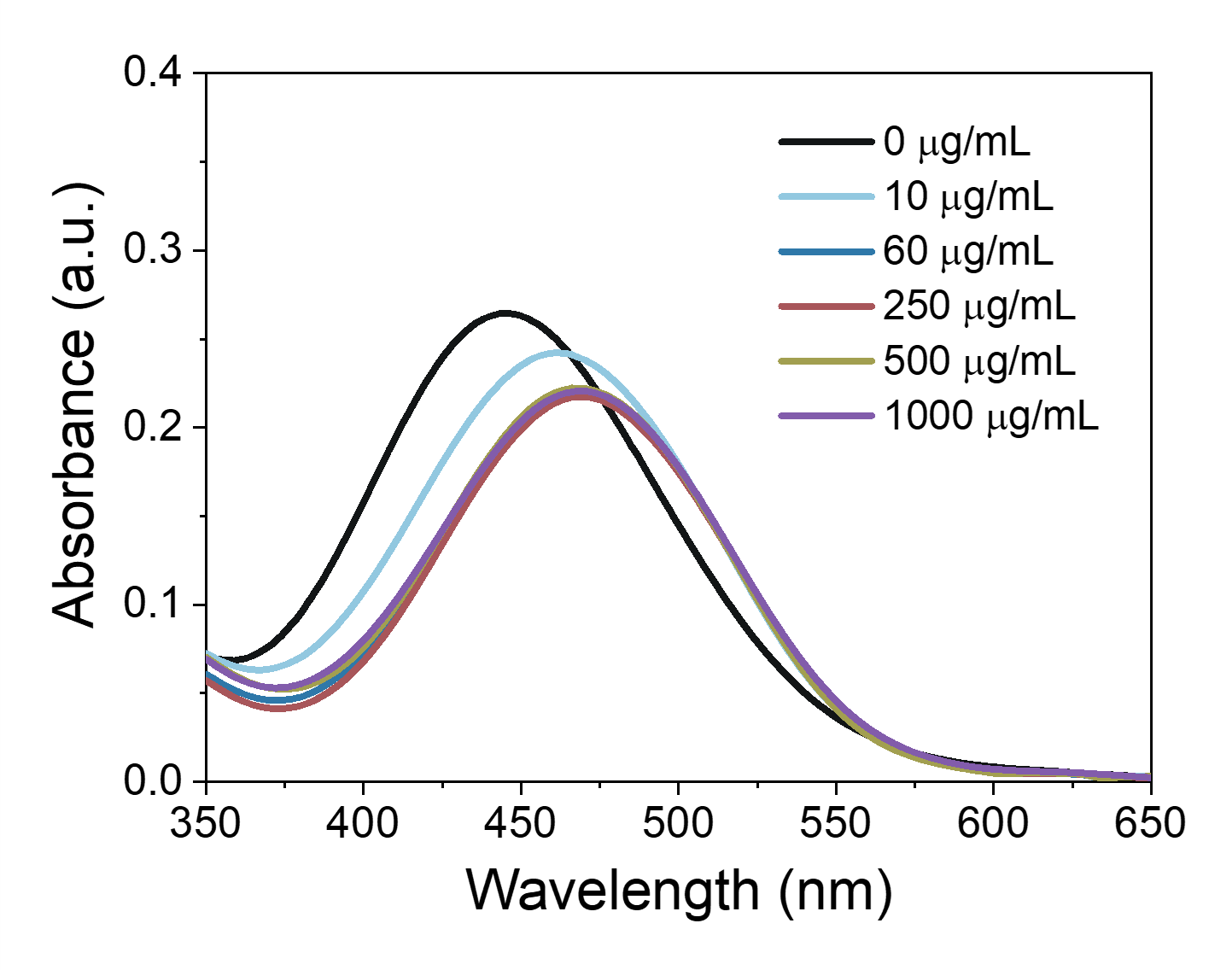

**Supplementary Fig. 15** Absorption spectra of PVC-1 in buffer solutions with adding different concentrations of calf thymus DNA.

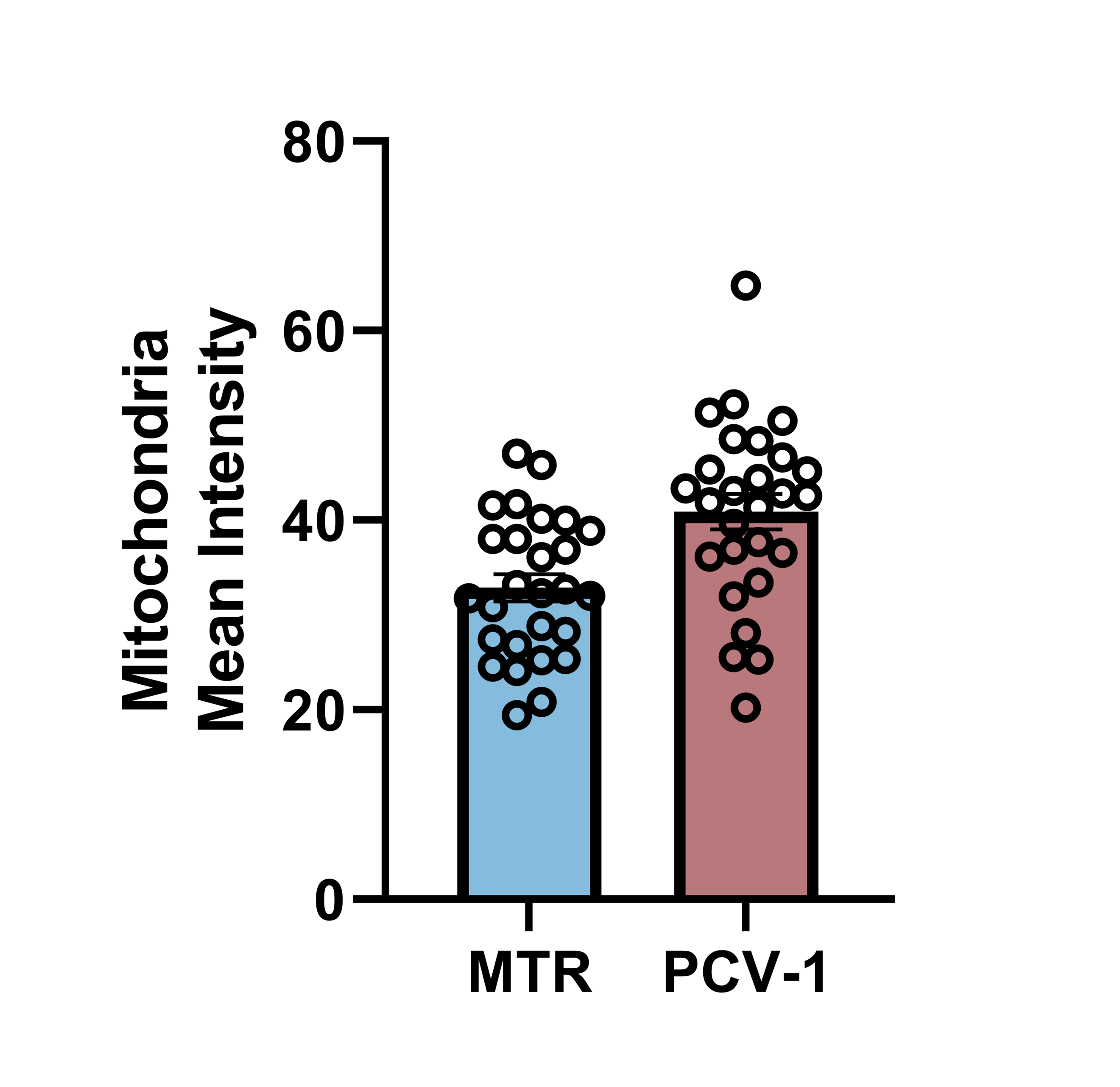

**Supplementary Fig. 16** Mitochondria mean intensity of MTR and PCV-1. Data are given as *M ± SEM*, *n* = 27.

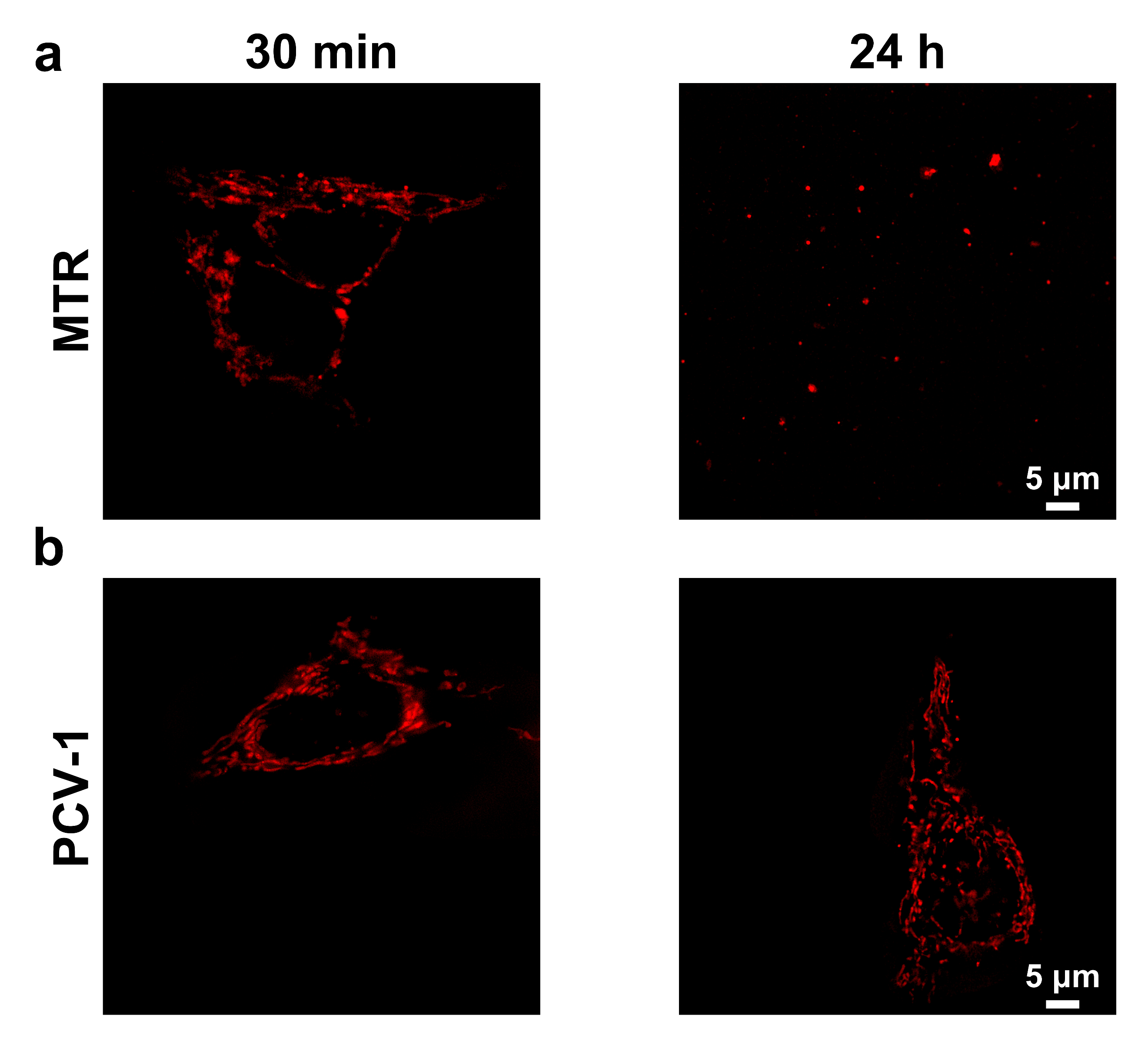

**Supplementary Fig. 17** **a** SIM images of HeLa cells stained with MTR for 30 min and 24 h. **b** SIM images of HeLa cells stained with PCV-1 for 30 min and 24 h.

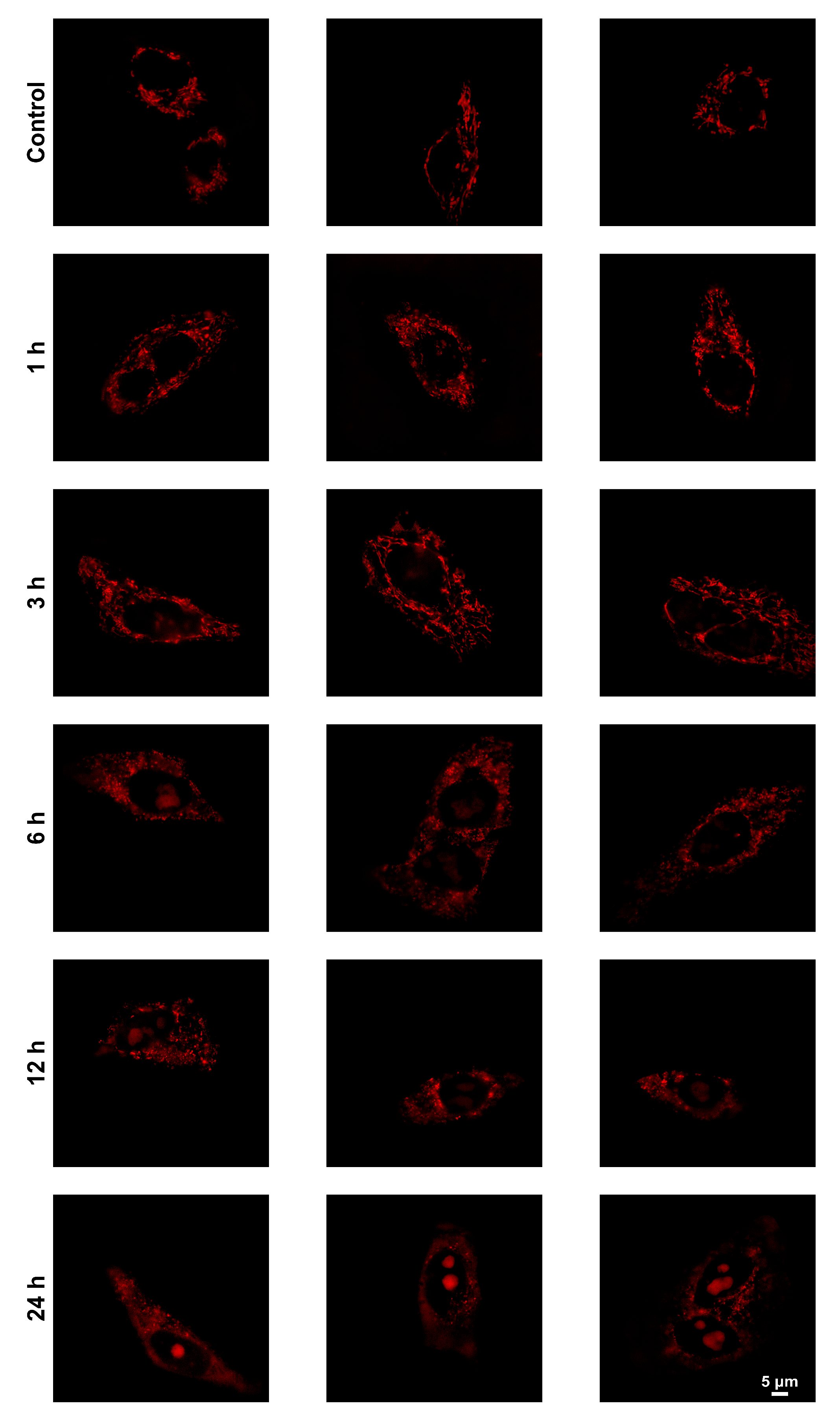

**Supplementary Fig. 18** The data set for **Figure 6d**.

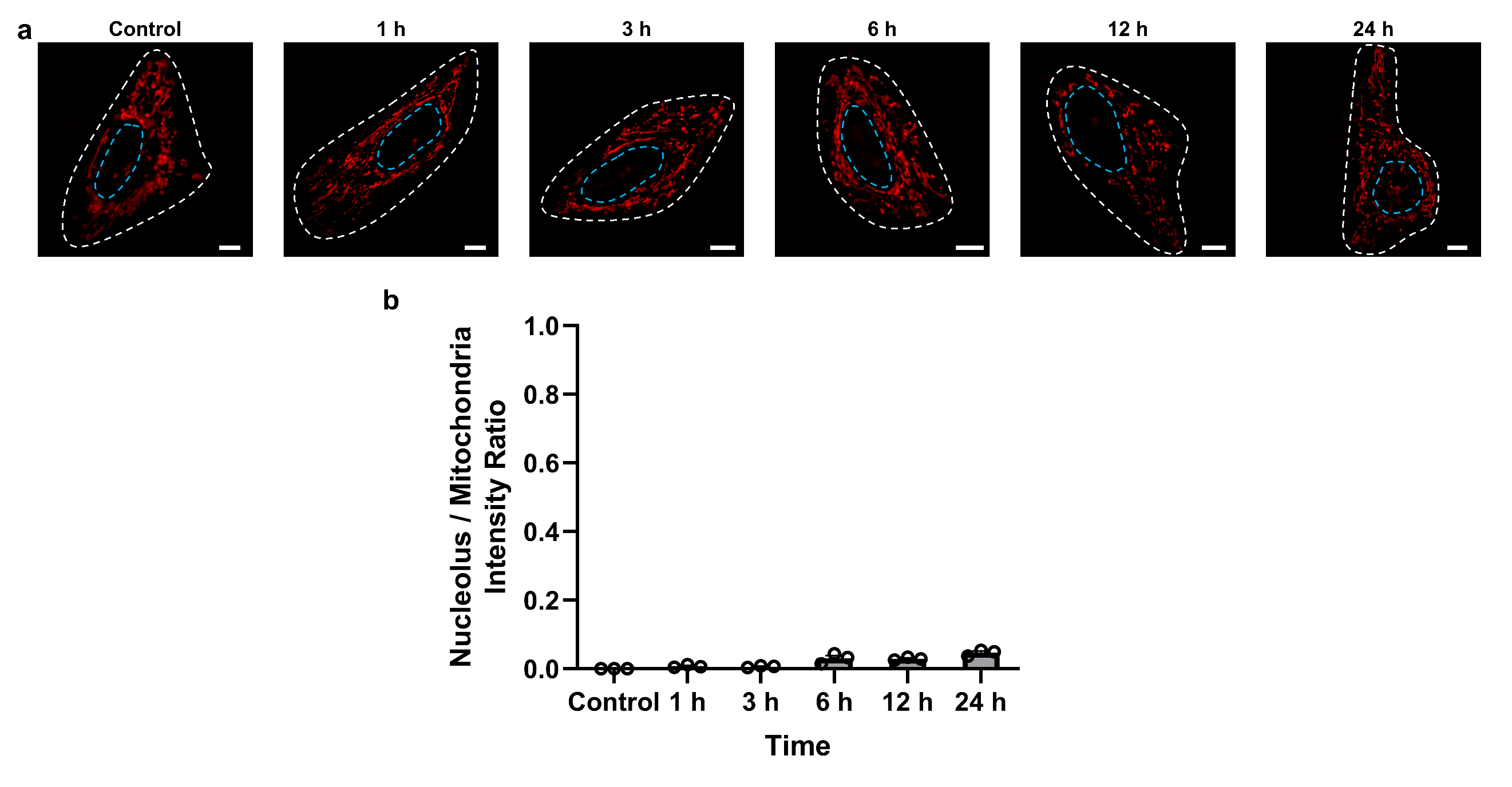
 **Supplementary Fig. 19** **a** SIM images of HeLa cells stained with PCV-1 at different times (0-24 h). PCV-1 channel: Ex 561 nm, Em 570-640 nm; scale bars: 5 μm. **b** The fluorescence intensity ratio of nucleolus to mitochondria in PCV-1-stained HeLa cells at different times (0-24 h). Data are given as *M ± SEM*, *n* = 3.

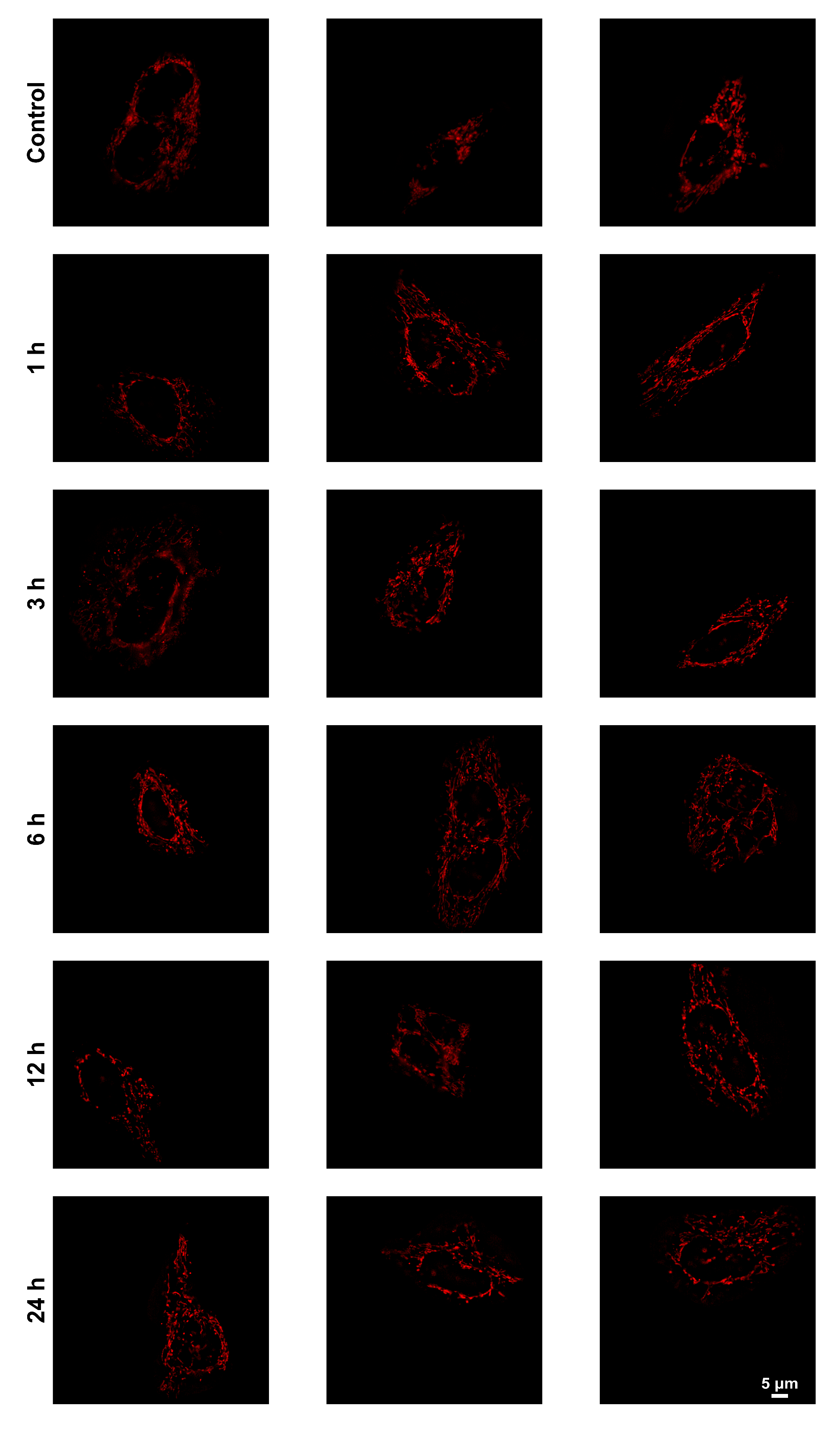

**Supplementary Fig. 20** The data set for **Supplementary Fig. 19 b**.

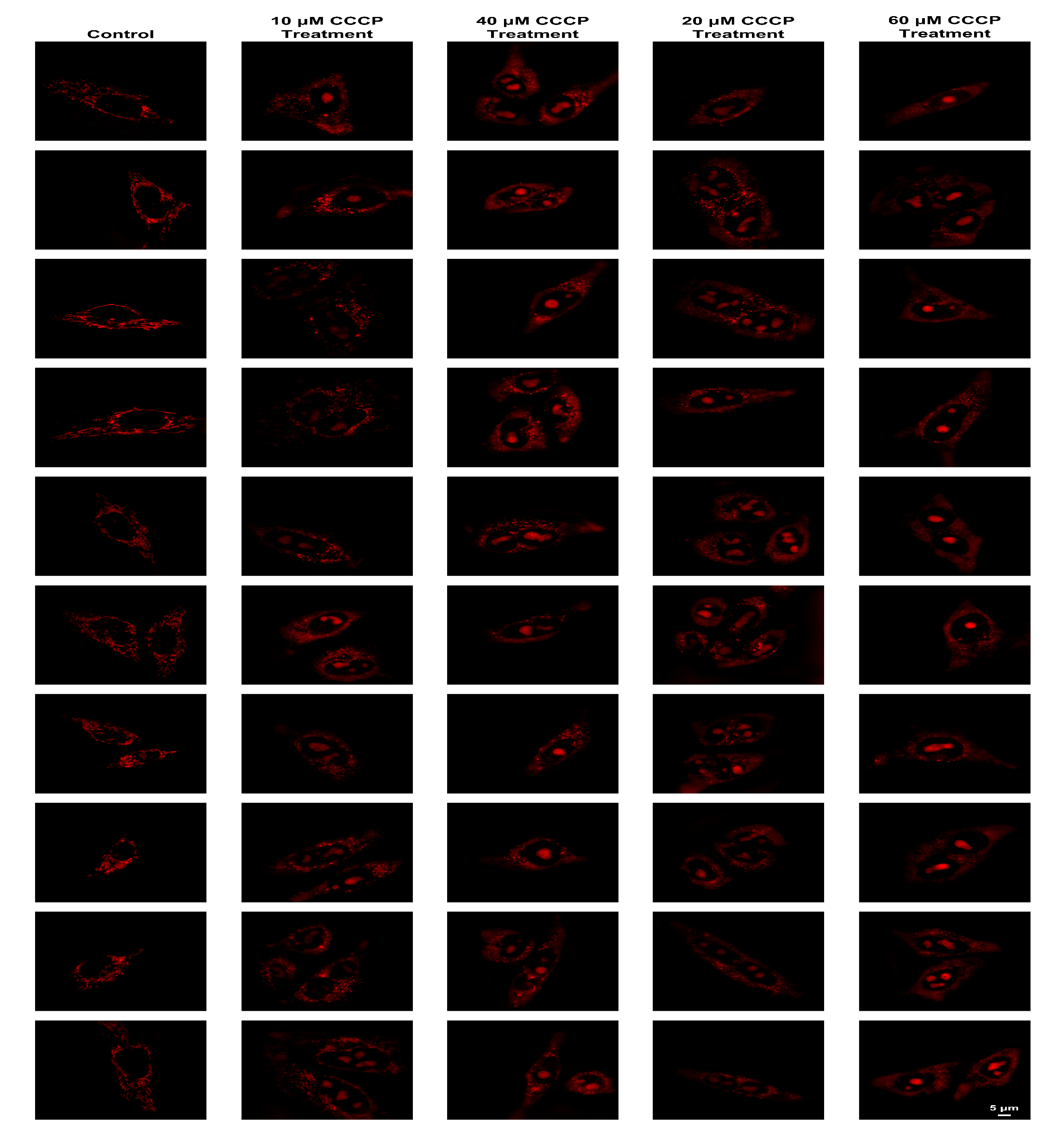

**Supplementary Fig. 21** The data set for **Fig. 7c**.

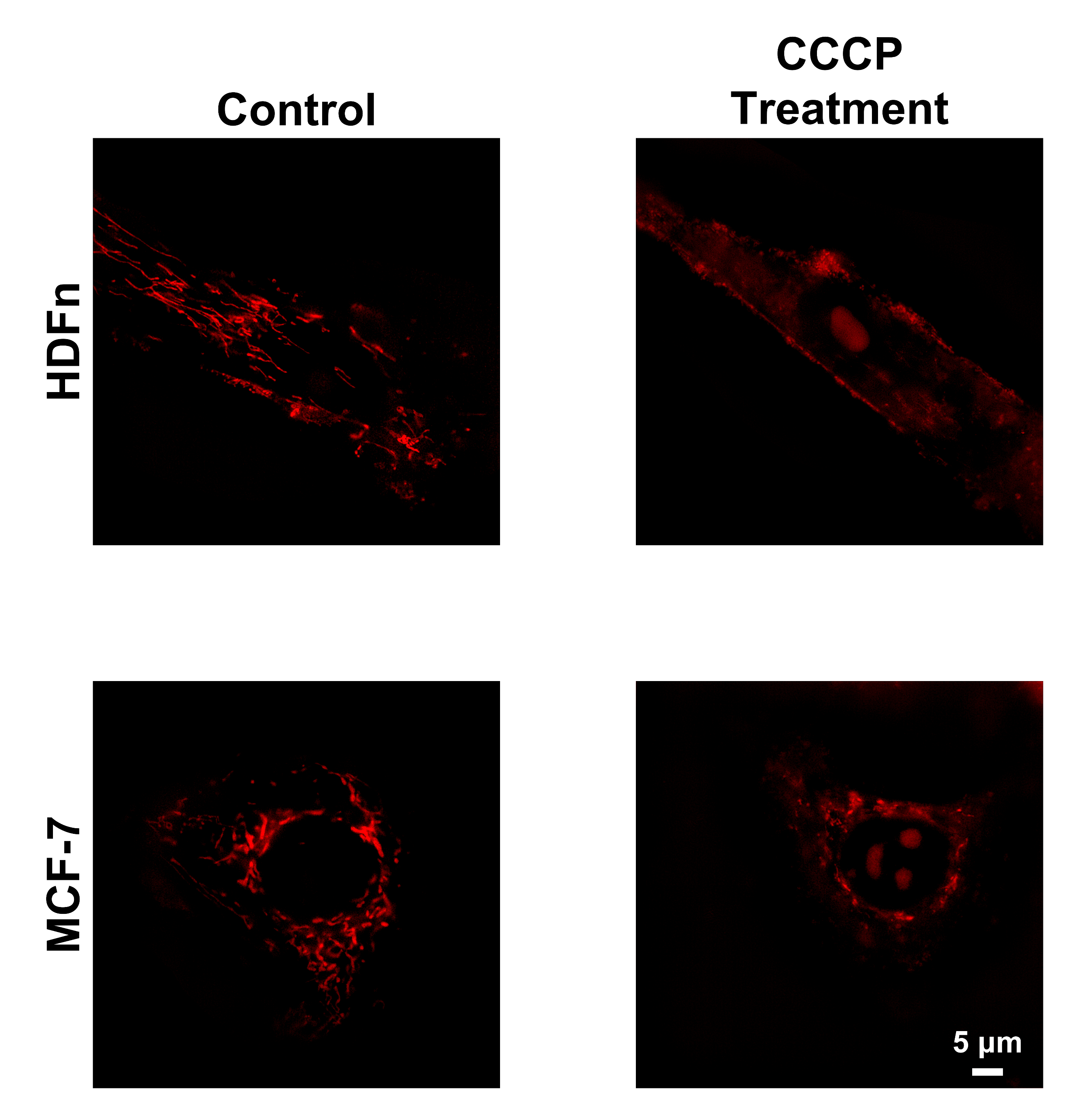

**Supplementary Fig. 22** SIM images of HDFn and MCF-7 cells treated with/without CCCP. CCCP treatment condition: 20 μM CCCP for 24 h.

**Supplementary Table 1** Calculated molecular orbitals of PCV-1 in H_2_O.

| MOs | Energy (eV) | Orbitals |
| --- | --- | --- |
| HOMO-5 (107) | -7.256 | 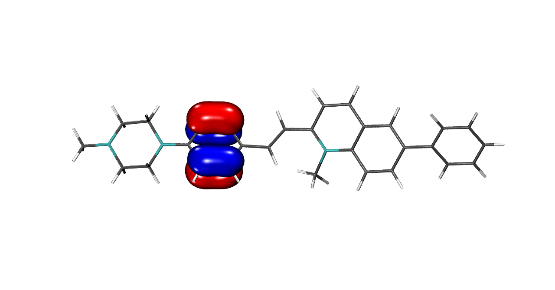 |
| HOMO-4 (108) | -7.113 | 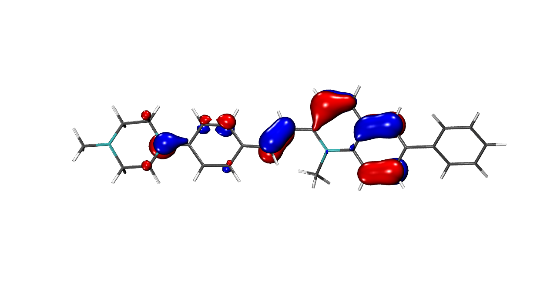 |
| HOMO-3 (109) | -7.039 | 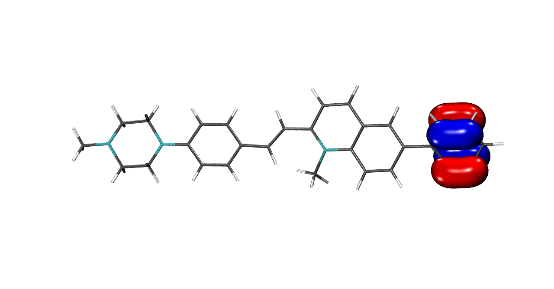 |
| HOMO-2 (110) | -6.509 | 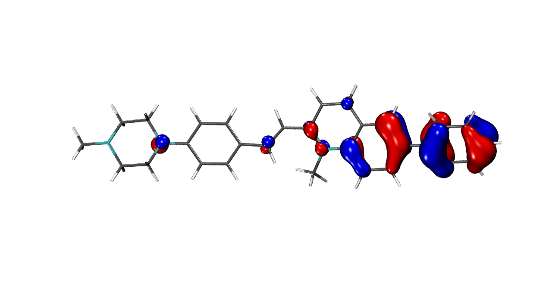 |
| HOMO-1 (111) | -6.086 | 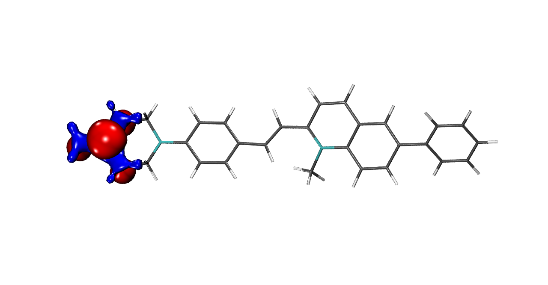 |
| **HOMO (112)** | **-5.473** | 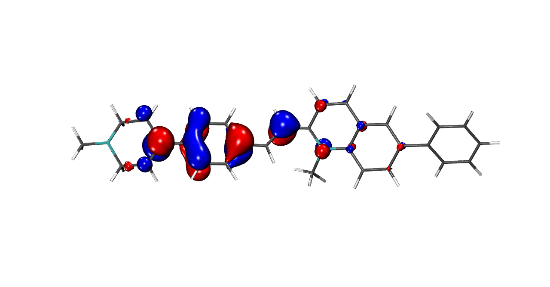 |
| **LUMO (113)** | **-2.947** | 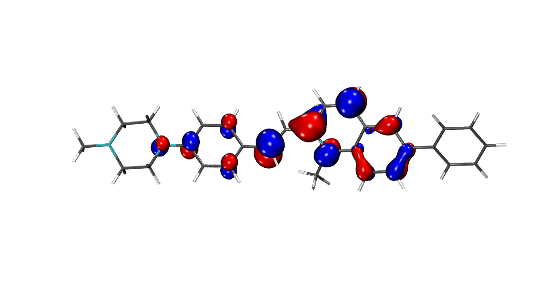 |
| LUMO+1 (114) | -1.612 | 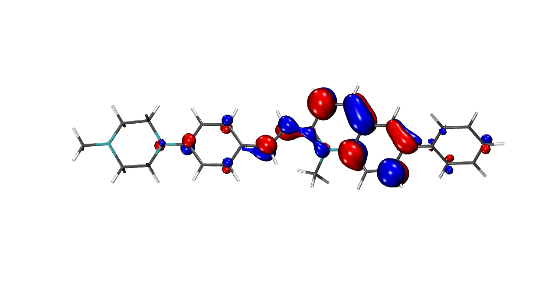 |
| LUMO+2 (115) | -1.086 | 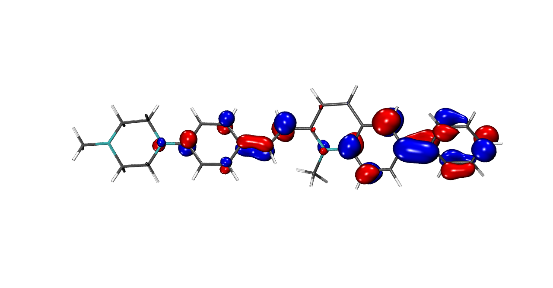 |
| LUMO+3 (116) | -0.343 | 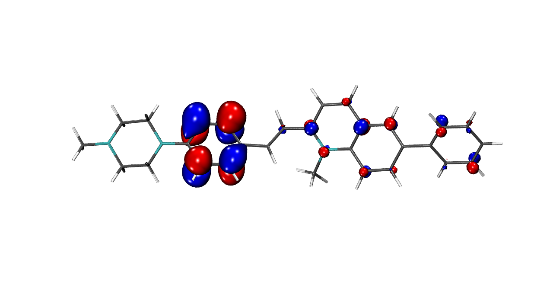 |
| LUMO+4 (117) | -0.278 | 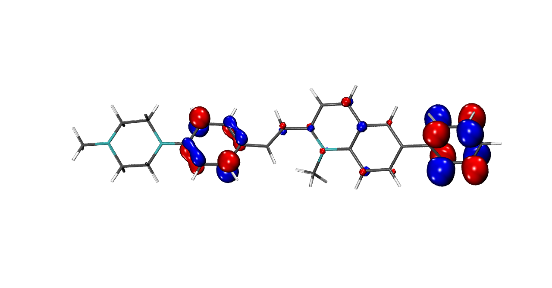 |
| LUMO+5 (118) | -0.182 | 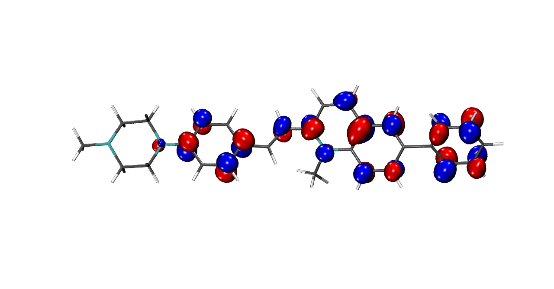 |

**Supplementary Table 2** Calculated singlet electron transitions (f > 0.01) of PCV-1 in H_2_O.

| No. | Wavelength (nm) | *f* | Major contributions |
| --- | --- | --- | --- |
| 1 | 533.4 | 1.4467 | HOMO->LUMO (100%) |
| 2 | 439.7 | 0.0186 | H-1->LUMO (100%) |
| 3 | 395.0 | 0.0573 | H-2->LUMO (94%) |
| 4 | 357.1 | 0.3432 | HOMO->L+1 (94%) |
| 5 | 338.0 | 0.0697 | H-4->LUMO (26%), H-3->LUMO (66%) |
| 6 | 335.7 | 0.0707 | H-4->LUMO (59%), H-3->LUMO (33%) |
| 8 | 307.0 | 0.0667 | H-6->LUMO (57%), HOMO->L+2 (33%) |
| 9 | 302.1 | 0.0157 | H-6->LUMO (35%), HOMO->L+2 (50%) |
| 10 | 294.8 | 0.0102 | H-1->L+1 (98%) |
| 11 | 280.1 | 0.3563 | H-2->L+1 (73%) |
| 12 | 276.0 | 0.0314 | HOMO->L+3 (72%), HOMO->L+4 (11%) |
| 15 | 256.9 | 0.0111 | H-3->L+1 (36%), HOMO->L+4 (39%) |
| 16 | 254.5 | 0.0123 | H-3->L+1 (26%), HOMO->L+4 (14%), HOMO->L+5 (38%) |
| 18 | 247.4 | 0.1342 | H-4->L+1 (10%), H-2->L+2 (82%) |
| 19 | 239.0 | 0.1413 | H-6->L+1 (10%), H-5->L+1 (10%), H-4->L+1 (52%) |
| 20 | 237.3 | 0.0435 | H-5->L+1 (57%), H-1->L+3 (22%) |
| 21 | 236.2 | 0.0199 | H-3->L+1 (11%), H-3->L+2 (43%), H-2->L+4 (15%) |
| 22 | 235.5 | 0.0485 | H-6->L+1 (11%), H-1->L+3 (32%), HOMO->L+6 (24%) |
| 23 | 235.2 | 0.0418 | H-6->L+1 (21%), H-5->L+1 (18%), H-1->L+3 (27%), HOMO->L+6 (19%) |
| 24 | 230.5 | 0.1628 | H-6->L+1 (31%), HOMO->L+6 (42%) |
| 26 | 223.6 | 0.0142 | H-9->LUMO (11%), H-4->L+2 (30%), H-1->L+4 (15%), H-1->L+5 (11%) |
| 27 | 223.2 | 0.0497 | H-10->LUMO (31%), H-8->LUMO (41%) |
| 28 | 222.4 | 0.0787 | H-9->LUMO (57%), H-4->L+2 (12%) |
| 29 | 219.4 | 0.0212 | H-10->LUMO (48%), H-8->LUMO (38%) |

Cartesian coordinates of optimized PCV-1

Charge = +1, Multiplicity = 1

C	8.31137100	-0.80448500	-1.01450200

C	9.69361800	-0.83946300	-1.18530500

C	10.50622900	0.08313400	-0.52332800

C	9.92748900	1.04275900	0.30940000

C	8.54491700	1.08099300	0.47874600

C	7.71605700	0.15604100	-0.17896100

C	6.24659400	0.19250600	0.00710100

C	5.48229700	-0.96859700	0.04401200

C	4.08361400	-0.93547700	0.21366200

C	3.42086200	0.31324300	0.36993200

C	4.18851700	1.49353800	0.32362400

C	5.55850200	1.42400500	0.14312900

C	3.29609200	-2.12076500	0.25028400

C	1.93905800	-2.04225900	0.35865800

C	1.26406200	-0.78923600	0.46086800

N	2.03362400	0.34281100	0.56530400

C	-0.16855700	-0.80896600	0.44442700

C	-1.03541300	0.16138800	-0.00173600

C	-2.45999400	0.07583300	-0.08639600

C	-3.18230200	1.13405500	-0.69056600

C	-4.55474400	1.11867700	-0.80898700

C	-5.32532700	0.02432900	-0.31880200

C	-4.59724100	-1.04661200	0.28746500

C	-3.22785100	-1.01425200	0.40037200

N	-6.69040300	0.00779200	-0.40015900

C	-7.46583300	1.15438600	-0.88897200

C	-8.78438700	1.28587800	-0.12025600

N	-9.55725100	0.05377200	-0.18746600

C	-8.79070000	-1.04162800	0.39101000

C	-7.48319800	-1.23065300	-0.37668600

C	-10.87215200	0.19449400	0.42728900

C	1.44753500	1.61170800	1.03474100

H	7.68704200	-1.50718100	-1.55951000

H	10.13568300	-1.58167100	-1.84367400

H	11.58366500	0.05544100	-0.65695900

H	10.55354500	1.75863600	0.83405000

H	8.10952700	1.81478200	1.15164000

H	5.96357600	-1.93849300	-0.03895000

H	3.72163600	2.46811800	0.38628000

H	6.11949500	2.35040800	0.07273400

H	3.78689700	-3.08580800	0.16190800

H	1.32711400	-2.93650400	0.33186200

H	-0.58002900	-1.78102900	0.69934000

H	-0.62074100	1.08914900	-0.38849200

H	-2.63466500	1.98248300	-1.09461100

H	-5.03513700	1.94360900	-1.31824800

H	-5.12299400	-1.89444000	0.70732700

H	-2.73634800	-1.84611100	0.89605600

H	-6.89648400	2.07177900	-0.74295600

H	-7.67249600	1.03544100	-1.96255200

H	-8.56154200	1.57949400	0.92546100

H	-9.36494900	2.09731900	-0.57321500

H	-9.37471700	-1.96565400	0.31448600

H	-8.56657200	-0.87494500	1.46405800

H	-7.71576800	-1.51200800	-1.41417900

H	-6.91148200	-2.04166000	0.06899400

H	-11.43100100	-0.73970500	0.31225900

H	-11.42981800	0.98863300	-0.07935700

H	-10.82350000	0.43890300	1.50497800

H	2.12879400	2.06323600	1.75704400

H	1.28204000	2.31013900	0.20921700

H	0.50274600	1.39955900	1.52993300
